# Microbiome-specific T follicular helper cells drive tertiary lymphoid structures and anti-tumor immunity against colorectal cancer

**DOI:** 10.1101/2021.01.27.428417

**Authors:** Abigail E. Overacre-Delgoffe, Anthony R. Cillo, Hannah J. Bumgarner, Ansen H.P. Burr, Justin T. Tometich, Amrita Bhattacharjee, Tullia C. Bruno, Dario A.A. Vignali, Timothy W. Hand

**Affiliations:** R.K. Mellon Institute for Pediatric Research, Pediatrics Department, Infectious Disease Section, UPMC Children’s Hospital of Pittsburgh, University of Pittsburgh, Pittsburgh PA, 15224.; Department of Immunology, University of Pittsburgh, School of Medicine, Pittsburgh PA, 15261.; Tumor Microenvironment Center, UPMC Hillman Cancer Center, Pittsburgh PA 15232.; Graduate Program of Microbiology and Immunology, University of Pittsburgh, School of Medicine, Pittsburgh PA 15213.

**Keywords:** Colorectal cancer, T follicular helper cell, microbiome, T cells, bacteria-specific, tertiary lymphoid structure, tumor microenvironment

## Abstract

Colorectal cancer (CRC) is a common and deadly disease, and patients with metastatic tumors often fail to respond to therapy. While select members of the microbiome are associated with improved anti-tumor immunity, mechanistic understanding of how the microbiome provides a benefit is lacking. We show that modification of the CRC-associated microbiome with a single immunogenic commensal bacteria can alter T cell differentiation, inhibit tumor growth, and increase survival. Microbiome-driven control of CRC required the formation of colonic tertiary lymphoid structures (TLS) and increased infiltration of the tumor with cytotoxic immune cells. In the context of CRC, CD4^+^ T cells specific to the newly introduced commensals differentiated into T follicular helper cells and were necessary for the formation of TLS, immune infiltration of the tumor, and control over CRC. Thus, modification of the intestinal T cell response by the microbiome can be used to augment anti-tumor immunity in colorectal cancer.

## Introduction

Colorectal cancer (CRC) is one of the most common and deadly forms of cancer, representing almost 10% of all cancer deaths worldwide. (Arnold et al., 2017; Bray et al., 2018; Favoriti et al., 2016; Rawla et al., 2018). Late stage metastatic CRC carries a very poor prognosis and is largely refractory to current treatments (Bray et al., 2018; Tauriello et al., 2017). Recently, immunotherapy has revolutionized the treatment of multiple kinds of epithelial cell cancers but has shown minimal effectiveness in CRC (Le et al., 2015). The reasons for this are not fully understood, highlighting a need for a deeper mechanistic understanding of the unique tumor microenvironment (TME) in CRC (Mlecnik et al., 2016).

The microbiome is a collection of microorganisms that live alongside or directly against barrier surfaces, and in mammals the largest population lives in the colon (Gilbert et al., 2018). The colonic microbiota has a large impact on multiple aspects of host biology by digesting and modifying diet and host-derived compounds. Specifically, the microbiota plays a critical role in shaping the host immune response and in particular, different members of the microbiota have distinct abilities to induce T cell activation and differentiation (Th1, Th17, T_FH_, T_reg_ etc.) (Atarashi et al., 2015; Honda and Littman, 2016; Lathrop et al., 2011). Immune responses to intestinal resident bacteria can be modified by the local environment, with infection, inflammation and diet all having important effects on the development and differentiation of microbiome-specific T cell responses (Ansaldo et al., 2019; Belkaid and Hand, 2014; Hand et al., 2012; Ivanov et al., 2009; Rothschild et al., 2018; Wegorzewska et al., 2019; Xu et al., 2018). Thus, each interaction between the immune response and a member of the microbiota is highly contextual. For instance, *Helicobacter hepaticus* (*Hhep*), an adherent bacteria that resides predominantly in the cecum and colon, induces local immune responses that vary broadly depending on the immune status of the host (Chai et al., 2017; Xu et al., 2018). Specifically, in conditions of health, *Hhep* colonization induces CD4^+^ T cells that differentiate into regulatory (T_reg_) or follicular helper (T_FH_) states but under immunodeficient settings, drives the differentiation of inflammatory *Hhep*-specific Th1 and Th17 T cells (Kullberg et al., 2003; Xu et al., 2018).

Members of the colonic microbiome can play a detrimental role in the health of the host and potentially contribute to diseases, including colorectal cancer. Indeed, some intestinal bacteria, such as pks+ *E. coli* and enterotoxigenic *B. fragilis*, can have a ‘pro-tumor’ effect by adhering to and directly interacting with colorectal tumors, driving mutagenesis and induction of inflammatory cytokines (Arthur et al., 2012; Dejea et al., 2018; Grivennikov et al., 2012; Kostic et al., 2013; Schwabe and Jobin, 2013; Sears and Garrett, 2014). Alternatively, some compositions of the microbiome and specific organisms are associated with immunotherapy responses in animal models and melanoma patients (Gopalakrishnan et al., 2018; Iida et al., 2013; Matson et al., 2018; Routy et al., 2018), highlighting the therapeutic potential of modifying the microbiome to modulate downstream immune responses. *Helicobacter* is an example of how the complex host/microbiome relationship can affect colorectal cancer, since under circumstances of immunodeficiency *Helicobacter spp.* supports the development CRC, while in lymphoreplete mice the presence of *Helicobacter* is associated with control over colitis associated cancer (Erdman et al., 2003; Maloy et al., 2005; Rosshart et al., 2017). Unfortunately, our ability to use the microbiome to augment tumor immunity is limited because the basic underlying mechanisms that determine whether a given commensal bacteria contributes to tumor growth or control are unknown. Further, while most anti-tumor immunotherapy relies on activating T cells, the role of microbiome-specific T cells in promoting or limiting immune responses against tumors is not well described. We hypothesized that innately immunogenic bacteria, such as *Hhep,* that inhabit the mucus layer and adhere to the colonic epithelium would be more likely to modulate the immune response and might have an effect that exceeds their relative low abundance (Fox et al., 1994; Li et al., 2015). Using a carcinogen-induced orthotopic mouse model of colorectal cancer, we have found that modification of the microbiome via *Hhep* colonization increases immune infiltration of the tumor, limits tumor burden and significantly extends overall survival. Further investigation revealed that the positive effects of *Hhep* colonization required the development of *Hhep*-specific CD4^+^ T follicular T helper cells (T_FH_) and the T_FH_ -dependent formation of tertiary lymphoid structures (TLS) within and around tumors. Therefore, bacteria-specific CD4^+^ T cell responses can contribute to anti-tumor immunity in CRC and may represent a new target to augment cancer immunotherapy.

## Results

### *H. hepaticus* remodels the tumor microenvironment and leads to long-term survival in colorectal cancerss

We hypothesized that a bacteria-specific immune response during tumor development may lead to enhanced anti-tumor immunity but that any effects will be highly dependent upon the properties of the bacteria chosen. Therefore, we sought to rationally modify the colonic microbiome through the addition of an intrinsically immunogenic intestinal bacteria: *Helicobacter hepaticus* (*Hhep*) (Danne and Powrie, 2018; Kullberg et al., 2006; Xu et al., 2018). *Hhep* is an ideal candidate, since the anti-*Hhep* T cell immune response is both dependent upon the local environment and has also been shown to induce colitis. (Xu et al., 2018). Here, *Hhep* colonization of tumor-bearing mice would allow us to directly assess whether local tumor development modified the differentiation of bacteria-specific T cells and subsequently, whether these changes affected tumor progression. Additionally, C57BL/6 mice from Jackson labs are tested to be free of *Hhep* and have a relatively homogenous intestinal microbiome which makes them a well-controlled system for these experiments. We colonized mice undergoing the azoxymethane-dextran sodium sulfate (AOM-DSS) model of colitis-associated colon cancer with *Hhep*, after tumors had developed (week 7) and assessed weight gain, tumor burden, and long-term survival (**Fig. 1A and Fig. S1A**). AOM-DSS was an attractive model for these studies because it is responsive to the intestinal microbiome and is also characterized by slow developing colon tumors, wherein we can parse the local effects on the immune response during the various stages of tumor development. Strikingly, in contrast to *Il10*^−/−^ or T cell deficient mice, we found that *Hhep* colonization of tumor-bearing lymphoreplete C57BL/6 mice leads to a significant survival advantage (**Fig. 1B**), and a reduction in both tumor number and size (**Fig. 1C-D**) (Ge et al., 2017; Kullberg et al., 1998; Nagamine et al., 2008). The anti-tumor effect of *Hhep* was conserved when it was gavaged prior to the initiation of the protocol and a similar trend was observed even when *Hhep* was given as late as week 9 post-tumor induction (**Fig. S1B**). Together, these data suggest that *Hhep* colonization alone mediates a potent anti-tumor effect in the colon.

**Figure 1:**
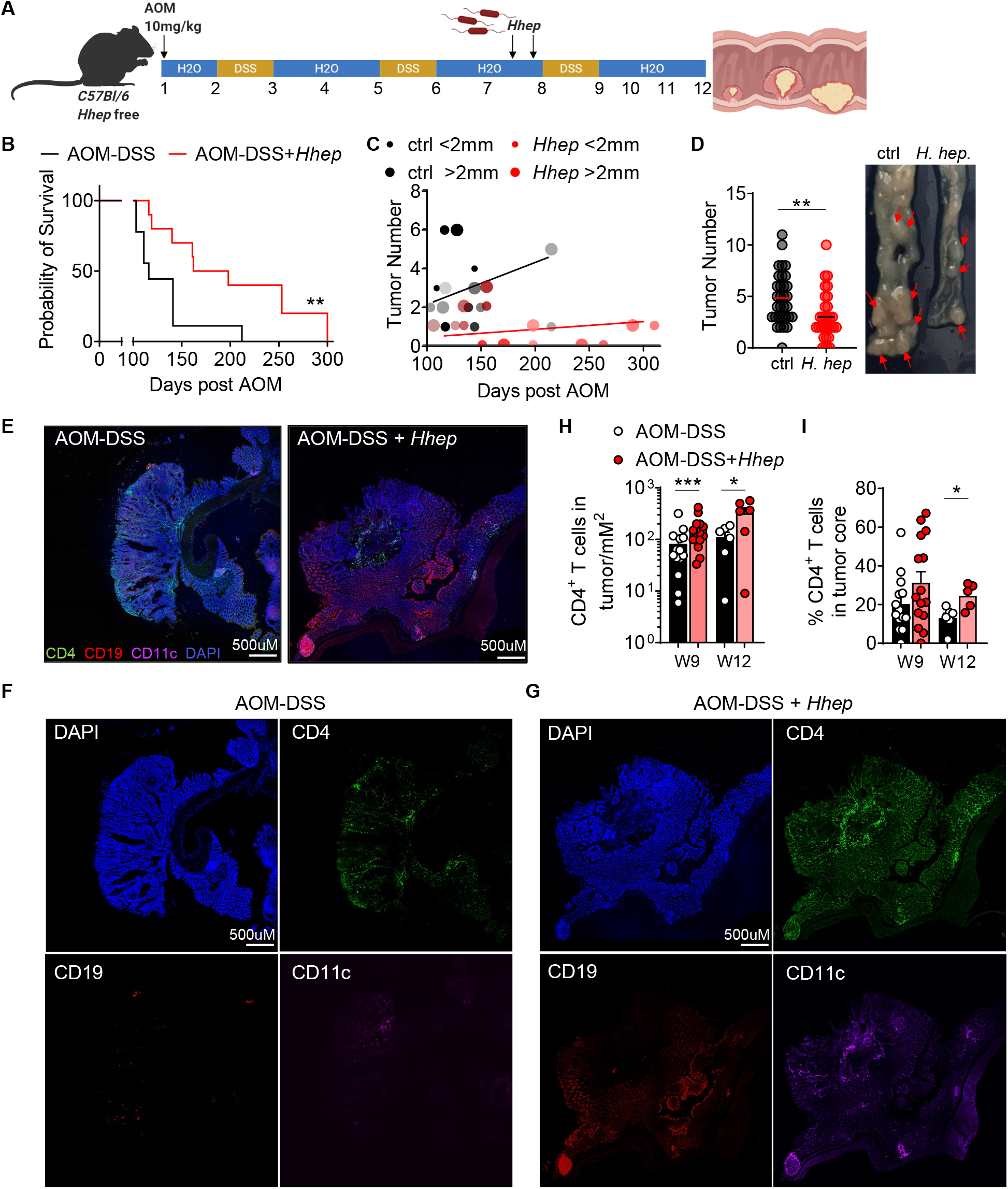
*H. hepaticus* remodels the tumor microenvironment and leads to long-term survival in colorectal cancer. *C57Bl/6 Hhep-*free mice were injected i.p. with 10mg/kg AOM on D0 and given 3% DSS in their drinking water on D7-14, 28-35, and 49-56. Half of the mice were gavaged on D45 and 49 with *Hhep.* (**A**) Experimental setup. (**B**) Mice were treated as stated above, and mice were removed from study when an unresolved prolapse persisted more than 2 days or when more than 30% weight loss occurred, and survival plots were generated. (**C**) Tumor number and size from (**B**). (**D**) Tumor number and representative image 12 weeks into the AOM-DSS protocol. (**E-G**) Representative image of immunofluorescence staining of colon sections 12 weeks post AOM. Shown: CD4 (green), CD19 (red), CD11c (purple), and DAPI (blue). (**H-I**) Quantification of CD4 T cells within the total tumor or percent found in the tumor core from (**E**). Data are a composite of 2 (**B-C**) independent experiments with 9-10 mice per group, 7 (**D**) independent experiments with 4-5 mice per group, and 2 independent experiments with 3 mice per group and 2-3 tumors per mouse quantified. Error bars represent the mean ± SEM. Kaplan-Meier (**C**), one way ANOVA (**H**) and student’s T test (**D, I**) were used. *p<0.05, **p<0.005, ***p<0.0005.

The colonic microbiome can vary during disease and is a critical regulator of both the growth of colorectal tumors and anti-tumor immune responses, so we sought to determine how intestinal colonization with *Hhep* during tumorigenesis impacted the microbiome. Surprisingly, *Hhep* colonization had little effect on the general structure of the microbiome of mice carrying colorectal tumors (**Fig. S1C-E**). At later timepoints (D82), *Hhep-*colonized mice could be discriminated from untreated controls on the basis of their microbiome, with *Hhep*-colonized mice grouping closer to the healthy pre-treatment (D0) microbiome configuration (**Fig. S1C**), but the differences were not associated with statistically significant changes (as determined by LEfSe analysis) in any intestinal taxa (**data not shown**) and may represent that *Hhep*-colonized mice have begun to control CRC.

We hypothesized that *Hhep* colonization was increasing immune cell infiltration of the tumor which is typical of tumors which respond better to therapy (Galon and Lanzi, 2020; Mlecnik et al., 2016; Sharma et al., 2017). Imaging of tumor sections revealed that *Hhep* colonization led to a significant increase in CD4^+^ T cells, B cells and CD11c^+^ dendritic cells into the tumor (**Fig. 1E-G**). Quantitative analysis of our imaging data revealed that tumors in *Hhep*-colonized mice were populated with significantly more CD4^+^ cells and that these cells tended to be located within the tumor core and much less likely to be confined to vessels running through the tumor (**Fig. 1E-G & S2A**). Flow cytometric analysis indicated that there was a significant decrease in the ratio of CD4^+^Foxp3^+^ T_regs_ to CD4^+^Foxp3^−^ T cells within in the tumor compartment, consistent with an increase in tumor-resident effector CD4^+^ T cells (T_eff_) (**Fig. S2B-C**). Altogether we have shown that colonizing mice with an immunogenic commensal organism can lead to substantial control over tumor growth that is associated with increased immune infiltration of the tumor.

### *Hhep*-driven anti-tumor immunity is associated with an increase in cytotoxic lymphocytes within colorectal tumors

To more comprehensively capture the effects of *Hhep* colonization on the TME, we performed single cell RNAseq (scRNAseq) on CD45^+^/MHCI^+^ cells from both the tumor-containing epithelial layer (EL) and the lamina propria (LP) of tumor bearing mice 12 weeks post tumor induction. DeteRministic Annealing Gaussian mixture mOdels for clusteriNg Single-Cell data (DRAGON) clustering analysis of the combined EL and LP sequencing identified 16 unique cellular clusters comprising the major cell types known to reside in the colon (**Fig. S2D**). Further clustering analysis of the EL scRNAseq data indicated that *Hhep* colonization induced an increase in a mixed group of cytotoxic lymphocytes (CTLs) (Cluster 1) and effector T cells (Cluster 5) within the intestinal epithelium and TME, consistent with the hypothesis that *Hhep* colonization is promoting more effective anti-tumor immunity in CRC (**Fig. 2A-C**). Deeper analysis of scRNAseq data revealed that Cluster 1 contained a variety of different Natural Killer (NK) cells and T cells (CD8^+^, CD4^+^, NK) expressing genes related to cytotoxic function (Gzma, Fasl, Tbx21) (**Fig. 2D-E and S2E**). In accord, *Hhep* colonization also increased cellular expression of chemokine Cxcl10, associated with the trafficking of cytotoxic T and NK cells and the number of cells expressing its receptor Cxcr3 (**Fig. 2F and S2F**). Confocal microscopy revealed that *Hhep* colonization significantly increased cytotoxic lymphocyte accumulation (NK and T cells) in colorectal tumors (**Fig. S2G-H**). While CD8^+^ T cells are the most studied cytotoxic lymphocytes, both NK cells and CD4^+^ T cells have also been shown to be potent anti-tumor immune cells in a variety of different tumor types (Bihl et al., 2010; Doorduijn et al., 2017; Ma et al., 2018). Thus, *Hhep* colonization leads to increased tumor infiltration of CTLs, which support anti-tumor immunity and are associated with better responses to immunotherapy (Mlecnik et al., 2016).

**Figure 2:**
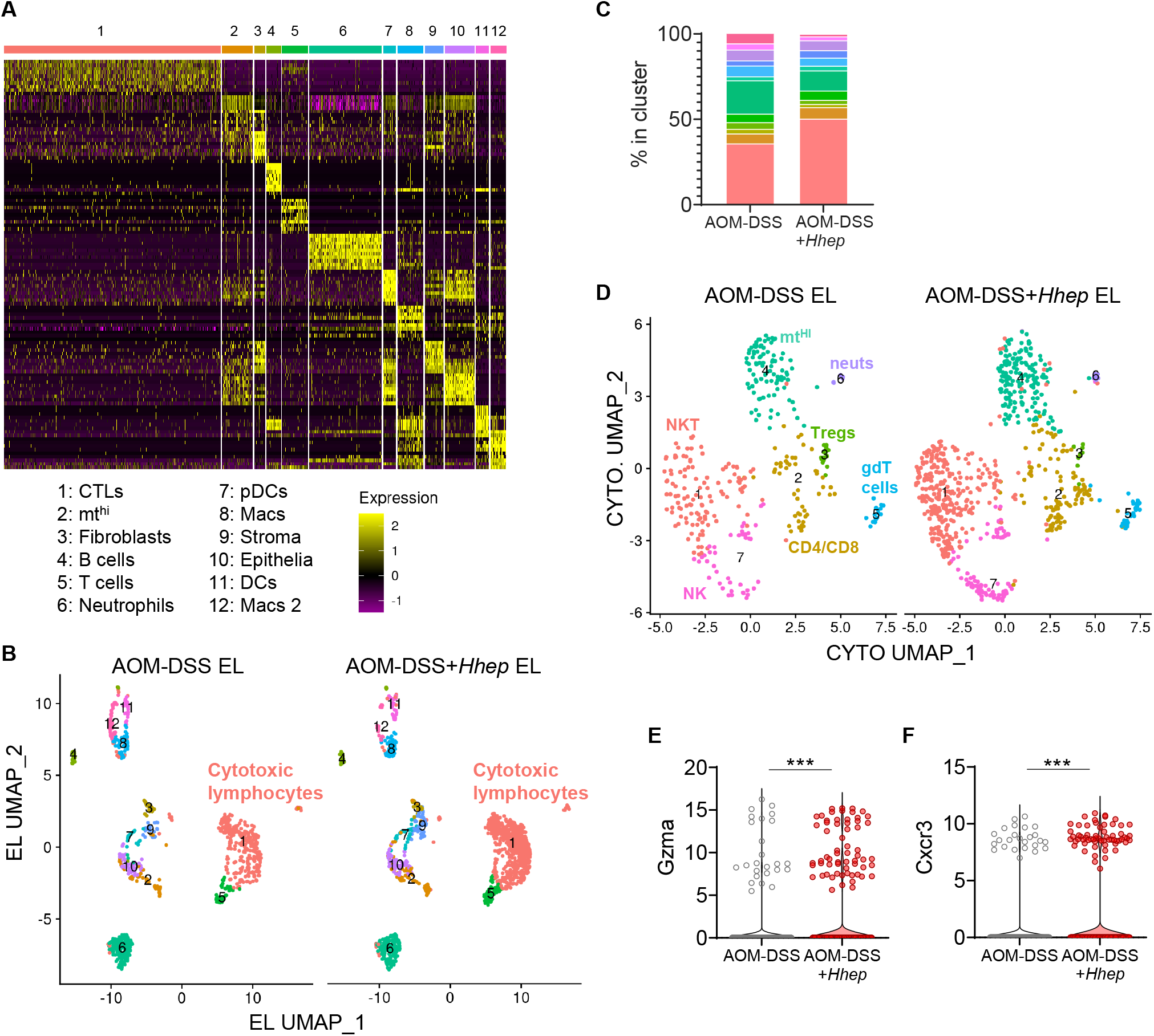
*Hhep-*driven anti-tumor immunity is associated with an increase in cytotoxic lymphocytes in and around the tumor. *Hhep* induces cytotoxic lymphocytes in and around the tumor. *C57Bl/6 Hhep-*free mice were injected i.p. with 10mg/kg AOM on D0 and given 3% DSS in their drinking water on D7-14, 28-35, and 49-56. Half of the mice were gavaged on D45 and 49 with *Hhep.* (**A-D**) Cells from LP and EL at week 12 of the AOM-DSS protocol (3 AOM-DSS mice, 3 AOM-DSS + *Hhep* mice) were enriched for CD45^+^ cells, labeled with CD45/MHCi cell hashing antibodies, and subjected to scRNAseq. (**A**) Heatmap of 12 unique clusters found within the EL samples. (**B**) UMAP visualization and DRAGON clustering of EL revealed 12 unique clusters across all samples. (**C**) Quantification of (**B**). (**D**) Subclustering of Cluster 1 (Cytotoxic lymphocytes; CTLs). 7 unique clusters were identified. (**E-F**) Violin plots of various genes found within the EL of AOM-DSS+/− Hhep mice. Data represent 1 independent experiment with 3-4 mice per group per experiment. Wilcoxon rank sum test was used **(E-F**). ***p<0.0005.

### Colonization drives *Hhep*-specific CD4^+^ T follicular helper cell expansion

The majority of immunotherapeutic approaches, including those in CRC, target CD8^+^ T cells, primarily through boosting cytotoxic effector function (Diaz and Le, 2015; Le et al., 2017; McLane et al., 2019; Twyman-Saint Victor et al., 2015). Underscoring the diversity of cell types that express cytotoxic functions found in colorectal tumors, *Hhep*-mediated anti-tumor immunity was independent of CD8^+^ T cells but was completely dependent upon the presence of CD4^+^ T cells (**Fig. 3A and S3A**). Therefore, we analyzed the colonic CD4^+^ T cell response to *Hhep* colonization. *Hhep* is known to induce both T_reg_ and T_FH_ CD4^+^ T cell responses, but how the anti-*Hhep* T cell response is shaped by CRC is not known (Chai et al., 2017; Xu et al., 2018). Flow cytometry of the colon LP revealed a durable, long-lived increase in both the frequency and number of T_FH_ (CXCR5^+^PD1^+^CD4^+^ T cells) after *Hhep* colonization (**Fig. 3C-D**). To analyze *Hhep-*specific CD4^+^ T cells, we utilized both MHC class II tetramers and a congenic CD45.1^+^ *Hhep* TCR transgenic mouse (HH5-1tg CD45.1) that allow for the measurement of *Hhep*-specific CD4^+^ T cells (**Fig. 3B-D and S3B-C**). Using these tools, the majority (>60%) of *Hhep*-specific CD4^+^ T cells developed into Bcl6, CXCR5 and PD1 expressing T_FH_ that persisted for at least 4 weeks post-*Hhep* colonization (**Fig. 3C-E and S3D**). This was in contrast to tumor-free controls (DSS + *Hhep*, without AOM), where most of the *Hhep*-specific T cells became T_regs_ (**Fig. S3E**) (Chai et al., 2017; Xu et al., 2018). In non-colonized mice, HH5-1 tg T cells could not be recovered from the colon, indicating that these cells require *Hhep* for activation and differentiation (**Fig. S3C**). Analysis of adoptively transferred T cells specific to a different *Hhep* epitope (HH7-2tg) in *Hhep* colonized mice also show that the majority were differentiating into T_FH_, indicating that this response is generalizable in the context of colorectal tumors (**Fig. S3F-I**). Collectively, our results support the hypothesis that in the context of colorectal tumors, *Hhep* colonization induces *Hhep*-specific T_FH_ cells in the LP.

**Figure 3:**
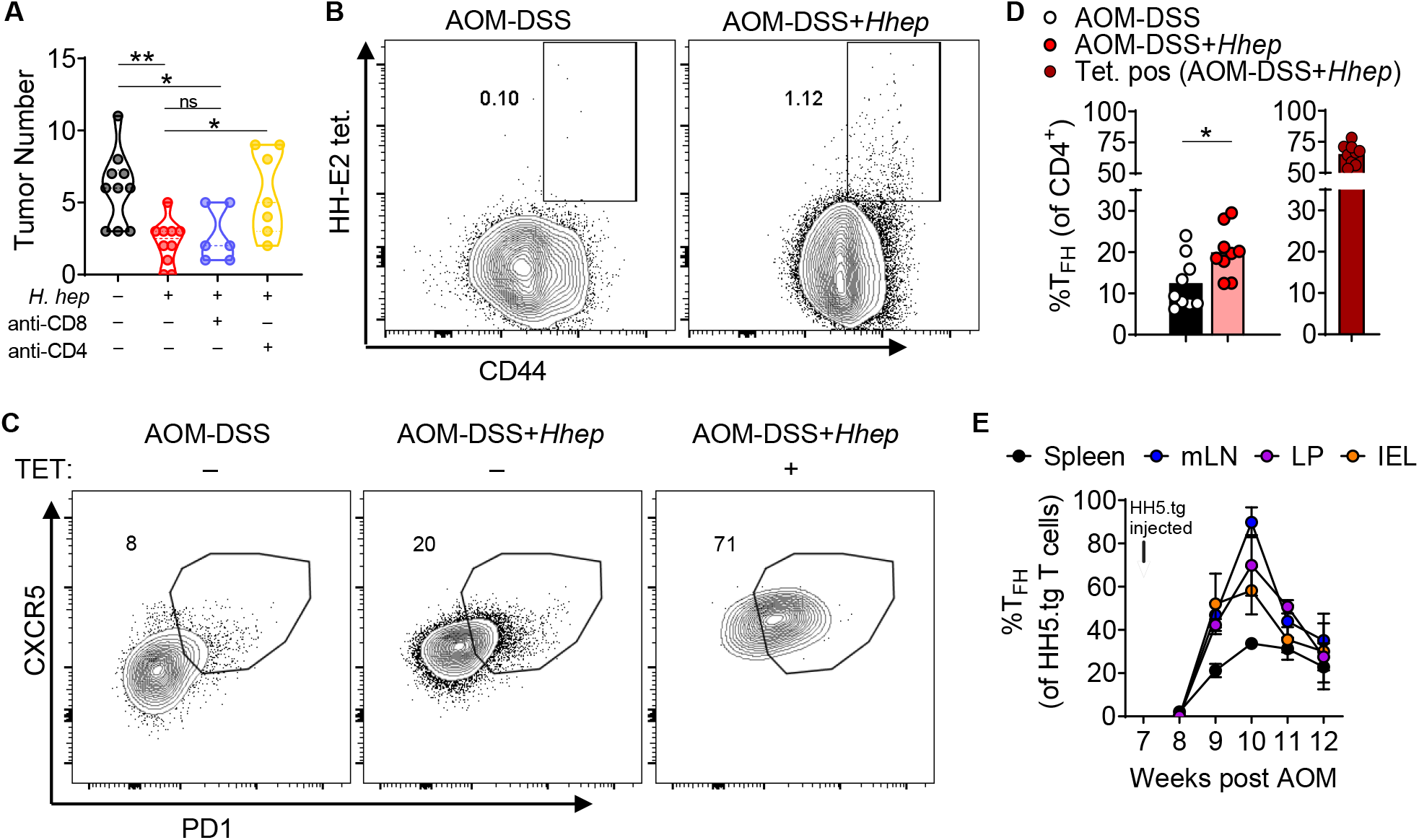
Colonization drives *Hhep-*specific T follicular helper cell expansion. (**A**) Tumor number quantification from mice given AOM-DSS +/− Hhep where half the mice were treated with anti-CD4 or anti-CD8 every 5 days starting on D58. (**B-E**) Cells were isolated from the colons of tumor-bearing mice +/− Hhep 12 weeks post AOM and enriched for Tetramer^+^ or CD45.1^+^ cells prior to staining. (**B**) Cells were isolated from the tumors of mice +/− *Hhep* 12 weeks post AOM and stained for flow cytometry. Shown are representative graphs of HH-E2 tetramer and CD44 to identify tetramer positive CD4 T cells from the LP. (**C**) Representative flow plot of tetramer positive and negative T_FH_ (gated on total CD4^+^ or Tetramer^+^ CD4^+^ T cells as indicated). LP cells were isolated from the colons of tumor-bearing mice +/− *Hhep* 12 weeks post AOM and enriched for *Hhep* tetramer^+^ cells prior to staining for flow cytometry. (**D**) Quantification of (**C**). (**E**) Time course of the percent of T_FH_ of transferred HH5-1tg CD4^+^ (*Hhep* TCR tg.) from different tissues, as indicated. Data represent 2-3 independent experiments with 3-4 mice per group per experiment. One-way ANOVA (**A**), and student’s T test (**D**) were used. *p<0.05, **p<0.005.

### *Hhep* colonization expands the colonic lymphatic network

B cells and CD4^+^ T cells predominantly accumulate in the colonic LP (**Fig. S4A**) (Chai et al., 2017; Xu et al., 2018). To comprehensively analyze the effect that *Hhep* and *Hhep-*specific T cells had on the LP we utilized scRNAseq (**Fig. S4B**). Unexpectedly, cluster analysis revealed an enrichment in *Hhep*-colonized mice for cells expressing genes associated with stromal and endothelial cells (clusters 1, 5, 9, and 13), and specifically an increase in a *Ccl21a^+^Lyve1^+^* lymphatics cluster (cluster 9, sub-cluster 2) (**Fig. 4A-B**). In accord with an expansion of the lymphatic network in the colon of *Hhep* colonized mice, we observed an increased frequency of genes required for lymphangiogenesis (*Pdpn, Vegfc*) in the LP and fluorescent microscopy revealed a significant increase in Lyve-1 stained vessel-like structures (**Fig. 4C-D & Fig. S4C-E**) (Chen et al., 2016; Farnsworth et al., 2019; Hunter et al., 2016; Martens et al., 2020). Lymphatic vessels express integrins and secrete cytokines that can recruit and support immune cells (T & B cells, DCs) potentially increasing lymphocyte traffic and anti-tumor immunity (Lund, 2016; Lund et al., 2016).

**Figure 4:**
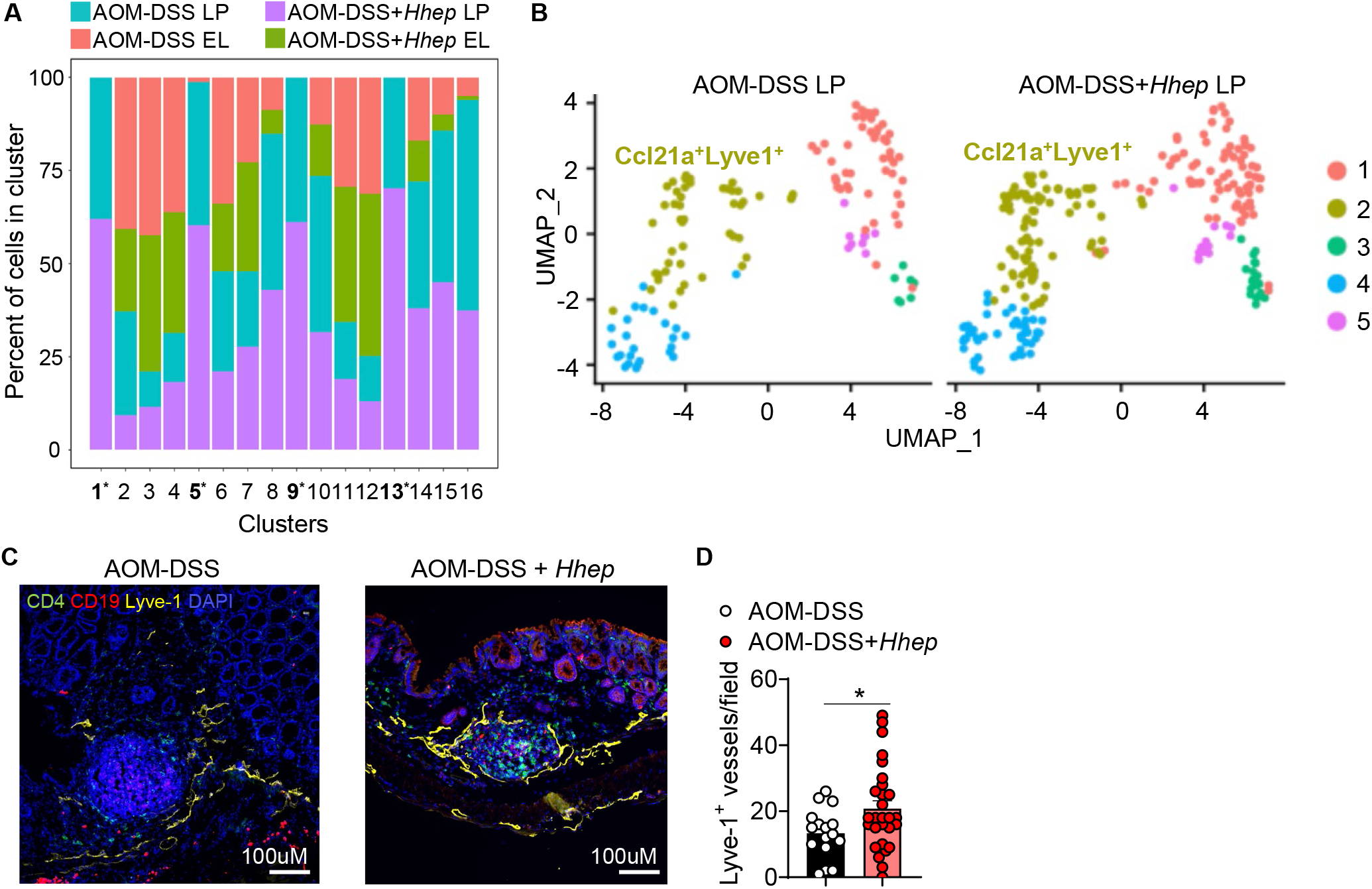
*Hhep* colonization expands the colonic lymphatic network. (**A**) Cells from the LP were prepared and sequenced as in (**Fig. 2**). DRAGON clustering revealed 16 unique clusters and an increase in the lymphatics cluster (cluster 9) in the LP of *Hhep*-colonized mice. (**B**) Subclustering of Cluster 9 (lymphatics) from scRNASeq LP samples. 5 clusters were identified, and Cluster 2 represents the Ccl21a^+^Lyve1^+^ cells. (**C**) Lyve1^+^ IF staining. Shown: CD4 (green), CD19 (red), Lyve1 (yellow), DAPI (blue). (**D**) Quantification of (**C**). Data represent 1 (**A-B**) or 3 (**C-D**) independent experiments with 3-4 mice per group. Student’s T test (**D**) was used. *p<0.05.

### Mature tertiary lymphoid structures form in response to *Hhep*

The expansion of lymphatic and stromal cells in the LP of *Hhep*-colonized may also be indicative of the development of tertiary lymphoid structures (TLS). TLS are ectopic lymphoid organs that form in response to chronic inflammation, and their presence inside or near tumors is associated with a positive prognosis for many tumor types including CRC (Becht et al., 2016; Di Caro et al., 2014; Coppola et al., 2011; Dieu-Nosjean et al., 2016; Lund, 2016; Lund et al., 2016; Petitprez et al., 2020; Ruddle, 2016). Indeed, *Hhep* colonized mice showed a significant increase in the number of organized TLS surrounding and within the tumor itself (**Fig. 5A-C & Fig. S5A-B**). Further, there was an obvious qualitative shift in the organization of TLS found in *Hhep* colonized mice, characterized by well-defined B and T cell zones and a significantly increased presence of CD11c+ cells in the T cell zone (**Fig. 5A**). TLS were observed as early as week 9 and were maintained at least through week 12 in the LP of *Hhep* colonized mice but were rarely seen in non-colonized mice (which predominantly contained relatively disorganized colonic patches), indicating that development and expansion of these structures requires *Hhep* and is not a generalized response to inflammation or tumor growth (**Fig. 5C and S5B**). In addition, the expression of genes associated with TLS formation were increased in the LP of *Hhep* colonized mice, such as *Lta, Tnfsf14* (LIGHT)*, Icam1,* and *Vcam1* (**Fig. S5C-F**). Thus, *Hhep* colonization is sufficient to drive an increase in lymphatics and TLS in the colonic LP that may be able to be better support T and B cell recruitment and the anti-tumor response.

**Figure 5:**
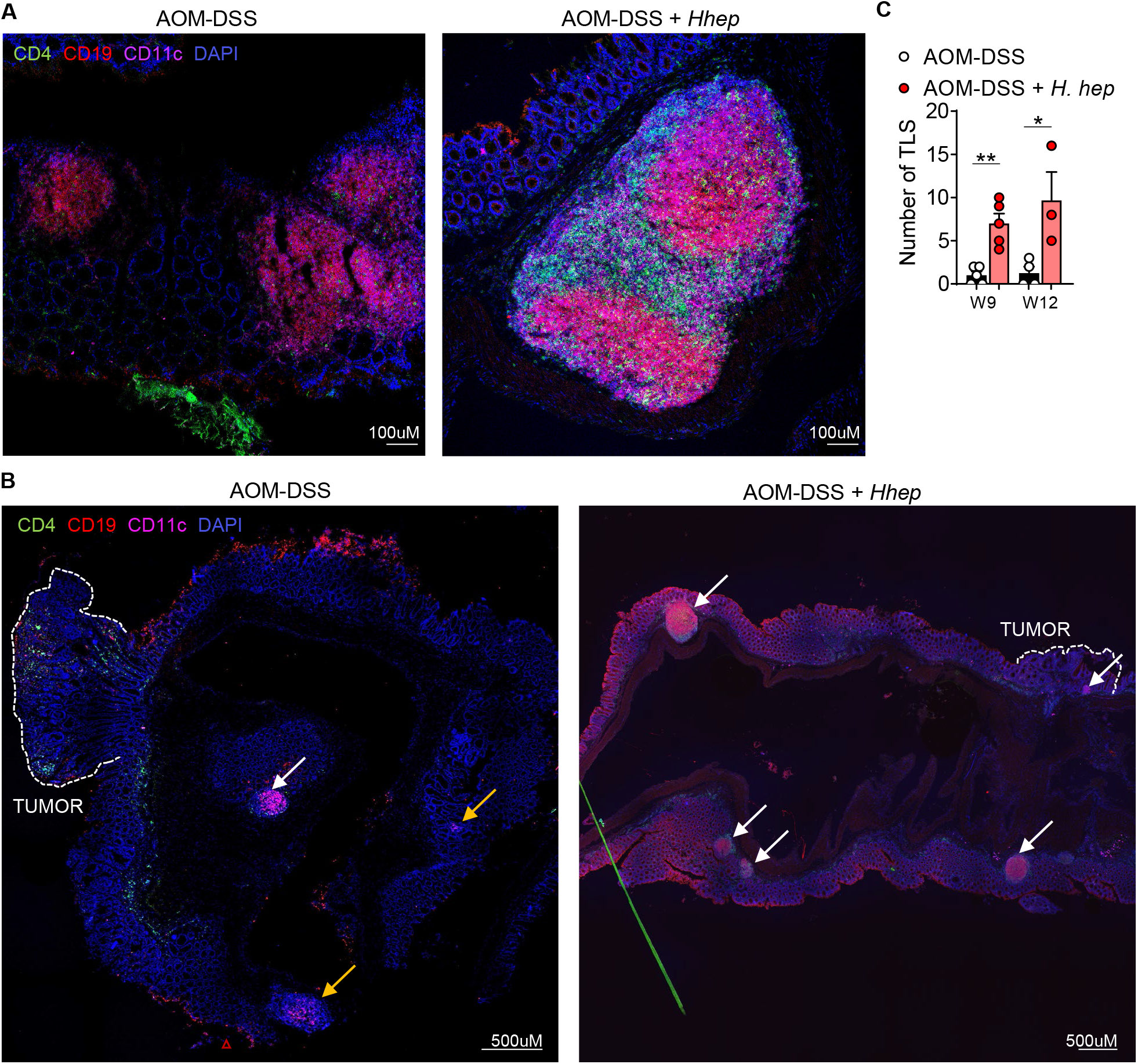
Mature tertiary lymphoid structures form in response to *Hhep*. *C57Bl/6 Hhep-*free mice were treated with AOM-DSS +/− *Hhep,* and colons were harvested 12 weeks post AOM and fixed with PFA prior to staining for IF. (**A**) IF staining of TLS. Shown: CD4 (green), CD19 (red), CD11c (purple), and DAPI (blue). (**B**) Large stitched images showing numbers and locations of TLS (white arrows) or colonic patches (orange arrows) found in AOM-DSS+/− *Hhep*. Shown: CD4 (green), CD19 (red), CD11c (purple), and DAPI (blue). (**C**) Quantification of number of TLS per colon of tumor-bearing mice +/− *Hhep.* Quantification performed at 9 and 12 weeks post AOM. Data represent 3-5 independent experiments with 3-5 mice per group. Student’s T test (**B**) was used. *p<0.05, **p<0.005.

### TLS contain *Hhep* and *Hhep*-specific T cells

*Hhep* is known to adhere to the colonic epithelium where it can be sampled by phagocytic immune cells, and we hypothesized that this immunogenicity may be important for the induction of TLS (Danne et al., 2017; Fox et al., 1994). To determine the biogeography of *Hhep* colonization, we utilized Fluorescent In-Situ Hybridization (FISH) for 16S rRNA genes. At both 9 and 12 weeks, we observed only sparse clusters of *Hhep* within the intestinal lumen (**Fig. S6A and data not shown**). Surprisingly, imaging of colon tissue of colonized mice consistently revealed bacteria, and specifically *Hhep* cells within nuclei dense regions of the lamina propria that are consistent with the structure of TLS (**Fig. 6A-C and S6B**). The presence of *Hhep* within TLS supports the idea that they may act as sites of activation for *Hhep*-specific T cells. To track the location of *Hhep*-specific T cells within the colon, we transferred naïve CD45.1^+^ *Hhep* TCR tg (HH5-1tg) CD4^+^ T cells into tumor-bearing WT mice 7 weeks post AOM administration. *Hhep*-specific CD4^+^ T cells were predominantly found within the TLS inside or around the B cell follicle (**Fig. 6D and S6C**). Neither *Hhep* (FISH analysis) nor *Hhep*-specific CD4^+^ T cells were consistently found within the tumor (**Data not shown**), supporting the notion that the CD4^+^ T cell-driven effects of *Hhep* colonization are dependent upon the CD4^+^ T_FH_ responses in the colonic LP, and more specifically tumor-associated TLS.

**Figure 6:**
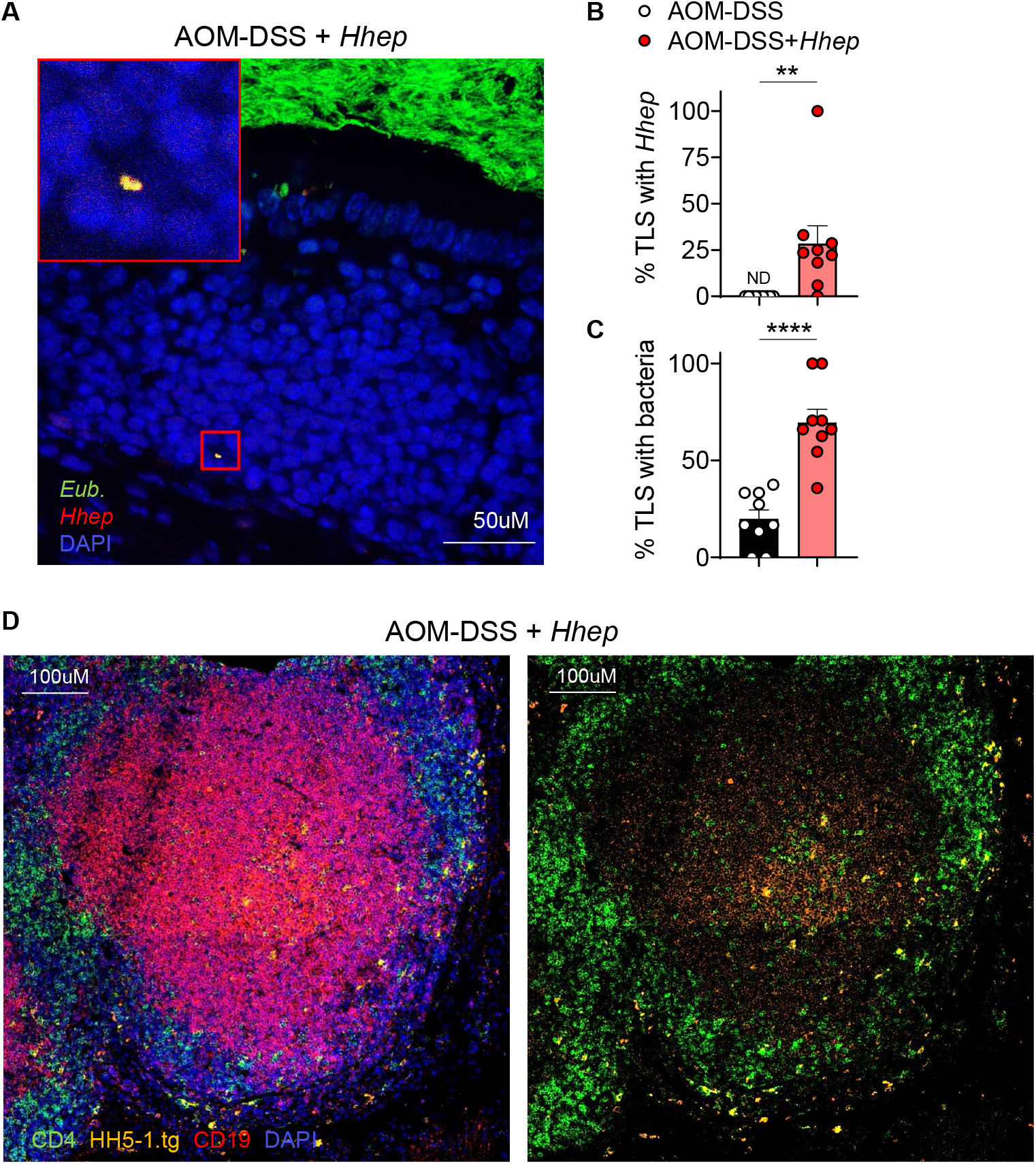
TLS contain *Hhep* and *Hhep*-specific T cells. *C57Bl/6 Hhep-*free mice were treated with AOM-DSS +/− *Hhep,* and colons were harvested 12 weeks post AOM and fixed with Methacarn (**A-C**) or PFA (**D**) prior to sectioning for IF. (**A**) FISH staining of TLS using 16S *Eubacteria* (green) and *Hhep* (red) specific probes. Yellow indicates co-staining. (**B-C**) Quantification of percent TLS containing *Hhep* or bacteria. (**D**) IF staining of TLS. Shown: CD4 (green), HH5-1tg CD45.1 T cells (orange), CD19 (red), and DAPI (blue). Data represent 3-5 independent experiments with 3-4 mice per group. Student’s T test (**B-C**) was used. **p<0.005, ****p<0.00005.

### *Hhep*-specific T_FH_ are necessary and sufficient to drive TLS formation and control tumor burden

T_FH_ cells are often found in and around B cell follicles, require B cells for development and survival, and provide B cell help through IL-21 secretion and CD40L expression (Crotty, 2019). One hypothesis consistent with our results is that the interaction of *Hhep*-specific CD4^+^ T_FH_ cells with B cells supports the growth of TLS in the colon. Accordingly, *Hhep*-mediated control over CRC required B cells (**Fig. S6D**) (Crotty, 2019; Helmink et al., 2020; Vinuesa et al., 2016; Yu et al., 2009). To test the importance of *Hhep*-specific T_FH_ in TLS induction and control over CRC directly, we utilized *Bcl6*^L/L^*Cd4*^Cre^ mice that largely lack T_FH_ (**Fig. 7A**) (Hollister et al., 2013; Johnston et al., 2009). AOM-DSS induction of CRC with *Hhep* confirmed that T_FH_ cells were required for *Hhep*-driven formation of TLS in the colon and control of tumor growth (**Fig. 7B-D**). Critically, transfer of *Hhep*-specific CD4^+^ TCR transgenic T cells into *Bcl6*^L/L^*Cd4*^Cre^ mice was sufficient to restore formation of organized TLS, increase immune infiltration into the tumor, and reduce tumor burden (**Fig. 7B-D and S7A-C**) (Cao et al., 2019). In aggregate, our data indicate that bacteria-specific T_FH_ are sufficient to drive TLS formation, increase immune infiltration, and lead to tumor reduction or clearance in an animal model of CRC. Both T_FH_ and intra- or peritumoral TLS have been associated with a positive response to therapy in head and neck cancer patients (Cillo et al., 2020) and CRC (Mlecnik et al., 2016; Schürch et al., 2020). Indeed, analysis of CRC patients within the TCGA database showed that an enriched T_FH_ signature generally presented within patients at a lower tumor stage and also associated with longer progression-free survival, while differences in CD8^+^ T cells showed no relationship with disease prognosis (**Fig. 7E-F & Fig. S7D**). Thus, both in mice and in human patients, the presence of T_FH_ and TLS is associated with better control over CRC and critically, within our animal model, microbiome-specific T_FH_ are necessary and sufficient to activate anti-CRC immunity.

**Figure 7:**
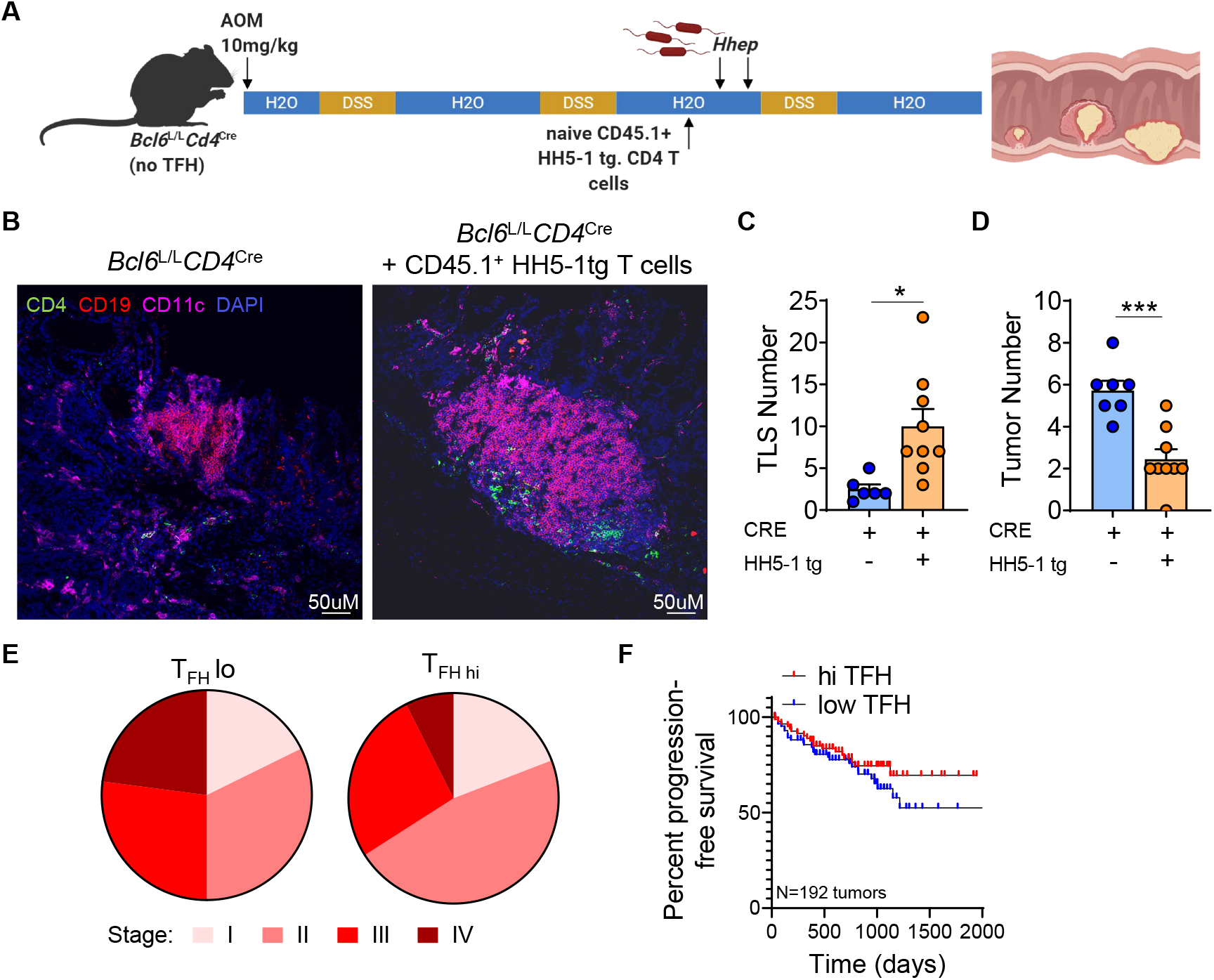
*Hhep-*specific T_FH_ cells are necessary and sufficient to drive TLS formation and control tumor burden. (**A**) Bcl6^L/L^*Cd4*^Cre^ mice were treated with AOM-DSS + *Hhep +/−* CD45.1^+^HH5-1tg T cells and tumor number and TLS numbers were assessed at week 12. (**B**) Colon sections were stained for CD4 (green), CD19 (red), CD11c (purple), and DAPI (blue) to assess TLS numbers. (**C-D**) Quantification of TLS (**C**) or tumor burden (**D**) of *Bcl6*^L/L^*Cd4*^Cre^ mice with or without HH5-1tg T cells transferred. (**E**) Tumor stage analysis of TCGA CRC patient data comparing those with a high or low T_FH_ signature. (**F**) Progression-Free Survival analysis of patients from (**E**). Data are a composite of 2 independent experiments with 3-5 mice per group (**B-D**). Error bars represent the mean ± SEM. Student’s t tests (**C-D**) was used. *p<0.05, ***p<0.0005.

## Discussion

The microbiome is an important modulator of the host immune response, and as such, particular members of the microbiota have been associated with positive prognosis in immunotherapy against melanoma. However, developing therapies based on promoting an anti-tumor microbiota has been challenging due to the fact that each study has identified distinct bacterial taxa (Gopalakrishnan et al., 2018; Matson et al., 2018; Routy et al., 2018; Sivan et al., 2015; Vétizou et al., 2015). Here, we present evidence that if the goal is to activate the immune response and foster anti-tumor immunity that efforts should be made to identify the most immunogenic bacteria, typically the bacteria that inhabit the mucus layer close to the intestinal epithelium (Ansaldo et al., 2019; Chiaranunt et al., 2018; Ivanov et al., 2009; Xu et al., 2018).

In our model, both control over tumors and immune infiltration of the tumor were associated with increased development of local TLS as well as the activation and differentiation of *Hhep*-specific T_FH_ cells. T_FH_ and the presence of mature TLS and have been associated with better prognosis of a variety of tumor types, but perhaps most notably CRC; however, the mechanism by which tumor-associated TLS activate anti-tumor immunity is not known (Becht et al., 2016; Di Caro et al., 2014; Cillo et al., 2020; Coppola et al., 2011; Dieu-Nosjean et al., 2016; Hollern et al., 2019; Lund, 2016; Lund et al., 2016; Petitprez et al., 2020; Ruddle, 2016). Here, we discovered that the invasion of the tumor by immune cells was dependent upon the presence of TLS. Why these structures are important to drive immune infiltration of the tumor is not known. We hypothesize that increased lymphatic density and TLS are leading to more efficient traffic of tumor-derived antigen and antigen-presenting cells, and thus are magnifying the local recruitment of tumor-specific T cells and the anti-tumor immune response. In addition, it is not currently known why some patients with CRC develop organized TLS while others do not. Here, we present evidence that for CRC the development of mature TLS is supported by the interaction of microbiota-specific T_FH_ with colonic B cells. We postulate that one reason for the heterogeneity in development of intra- and peri-tumoral TLS in different patients is differences in the intestinal microbiome, specifically in epithelial adherent bacteria known to induce T_FH_. We do not think that the induction of TLS and T_FH_ will be solely limited to *Hhep* as infection in the colon and other commensal bacteria can also induce these cells and structures (Ansaldo et al., 2019; Koscsó et al., 2020). The ability to induce TLS and tumor control may however be a property enriched in the *Helicobacter* genus. Indeed, gnotobiotic mice colonized with a *Helicobacter*-dominated microbiota from wild mice are partially protected from CRC induced by AOM-DSS (Rosshart et al., 2017). Conversely, some laboratory microbiomes containing other *Helicobacter* strains decrease the effectiveness of CD8^+^ T cell control over colorectal tumors (Yu et al., 2020), so it will be important to tease out the specific mechanisms employed by different strains of *Helicobacter* to induce mucosal immune responses. In humans, colonization with *Helicobacter pylori* has plummeted 2-3 fold over the past few decades with the lowest rates of infection being seen in young people (Jiang et al., 2012). In fact, loss of *H. pylori* has been used as a reference example for the ‘Vanishing Microbiota’ hypothesis which posits that microbiome diversity is narrowing over time with negative effects on human health (Blaser and Falkow, 2009). It would be interesting to learn if the prevalence of *Helicobacter* isolates that live in the colon is also declining. Indeed, it is possible that adding back immunogenic intestinal bacteria to the microbiome could be an important factor for the efficacy of current CRC patient treatments, including tumor immunotherapy (Asaoka et al., 2015; Diaz and Le, 2015; Topalian et al., 2012). Given we found that colonization led to an increase in T cell infiltration and function within the tumor, this may increase response rates to T cell targeted therapies such as anti-PD1. We propose that, in the future, it may be beneficial to combine rational targeted modification of the mucosal immune response by specific intestinal bacteria with current immunotherapeutic treatments that have so far been unsuccessful in the majority of CRC patients.

## Acknowledgements

The authors would like to thank A. Poholek for *Bcl6*^L/L^*Cd4*^Cre^ mice, J. Michel and A. Styche from the Children’s Hospital of Pittsburgh Flow Core for cell sorting, T. Tabib, R. Lafyatis and the UPMC Genome Center for preparation and sequencing of single cell RNA sequencing samples, the NIH tetramer core for MHCII tetramers containing sequences from *Helicobacter hepaticus,* the staff of the Division of Laboratory Animal Services for the animal husbandry, the Children’s Hospital of Pittsburgh Histology Core, the Center for Biological Imaging, the RK Mellon Institute, Department of Pediatrics, and the Department of Immunology at the University of Pittsburgh/UPMC as well as Nikhil Joshi, Grace Chen, Semir Beyaz, and the Hand Lab for helpful discussions. This work was supported by the UPMC Children’s Hospital of Pittsburgh/R.K. Mellon Institute for Pediatric Research, the NIH (R21 CA249074 to TWH; T32 5T32CA082084-18 to A.E.O.-D.; R01 CA203689 and P01 AI108545 to D.A.A.V.), the Damon Runyon Cancer Research Foundation (DRCRF postdoctoral fellowship to A.E.O.-D.), and the Hillman Cancer Center (Hillman Postdoctoral Fellowship for Innovative Cancer Research to A.R.C).

## Author Contributions

Conceptualization, A.E.O.-D. and T.W.H.; Formal Analysis, A.E.O.-D., A.R.C., H.A.B., A.H.P.B; Investigation, A.E.O.-D., H.A.B., A.H.P.B, J.T.T, A.B.; Resources, D.A.A.V. and T.W.H.; Writing— Original Draft, A.E.O.-D. and T.W.H.; Writing—Review & Editing, A.E.O.-D., A.R.C., H.A.B., A.H.P.B., J.T.T., A.B., T.C.B., D.A.A.V., and T.W.H.; Supervision, T.W.H.

## Declaration of Interests

DAAV: cofounder and stockholder – Novasenta and Tizona; stock holder - Oncorus and Werewolf; patents licensed and royalties - Astellas, BMS; scientific advisory board member - Tizona, Werewolf and F-Star; consultant - Astellas, BMS, Almirall; research funding - BMS, Astellas and Novasenta.

## Notes

### Competing Interest Statement

DAAV: cofounder and stockholder of Novasenta and Tizona; stock holder of Oncorus and Werewolf; patents licensed and royalties with Astellas, BMS; scientific advisory board member of Tizona, Werewolf and F-Star; consultant for Astellas, BMS, Almirall; research funding through BMS, Astellas and Novasenta.

