## Supplemental Files for "Microbiome-specific T follicular helper cells drive tertiary lymphoid structures and anti-tumor immunity against colorectal cancer"

Supplemental Figure 1: *Helicobacter hepaticus (Hhep)* reduces tumor burden but does not significantly alter the colonic microbiome.

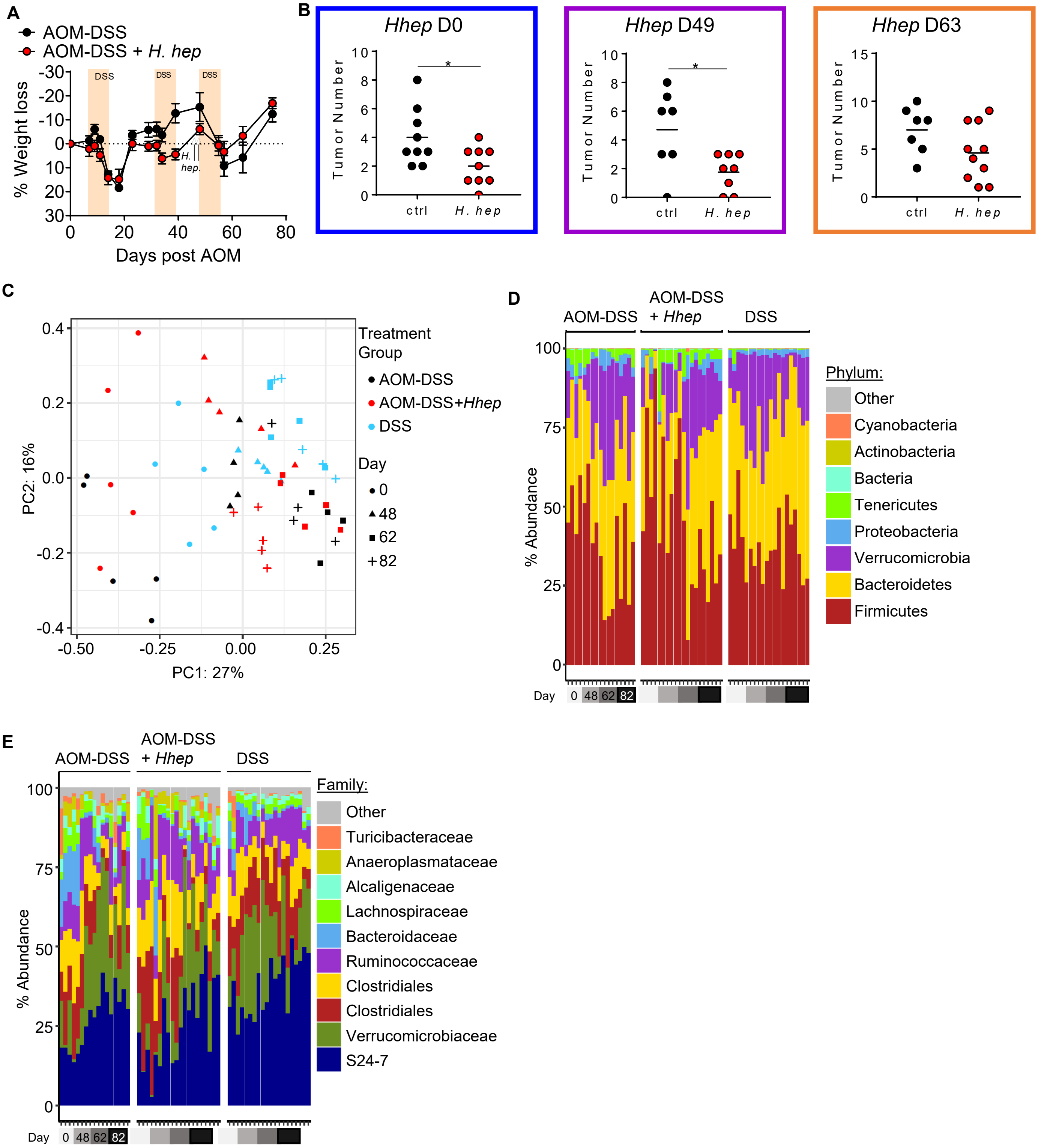

Supplemental Figure 1: *Helicobacter hepaticus (Hhep*) reduces tumor burden but does not significantly alter the colonic microbiome.

**Figure S1.**
(**A-B**) *C57Bl/6 Hhep*-free mice were injected i.p. with 10mg/kg AOM on D0 and given 3% DSS in their drinking water on D7-14, 28-35, and 49-56.
(**A**) Half of the mice were gavaged on D45 and 49 with *Hhep* and weights were monitored.
(**B**) Mice were treated as stated above and gavaged with *Hhep* on D0, D49, or D63 prior to tumor numbers being quantified at week 12 post AOM injection.
(**C-E**) Serial stool samples were taken from mice (as indicated) and DNA was isolated prior to 16S rRNA gene sequencing and analysis.
(**C**) Bray Curtis PCoA plot of 16S rRNA gene sequencing samples.
(**D-E**) Bar chart quantification of taxa represented in each group. (**D**) Phylum level and (**E**) family level. Data represent 2-5 (**A-B**) or 1 (**C-E**) experiment with 3-5 mice per group per experiment. Error bars represent the mean ± SEM. Student's T test (**B**) was used. *$p<0.05$.

Supplemental Figure 2: Colonization leads to an increase in cytotoxic lymphocytes within the epithelial layer.

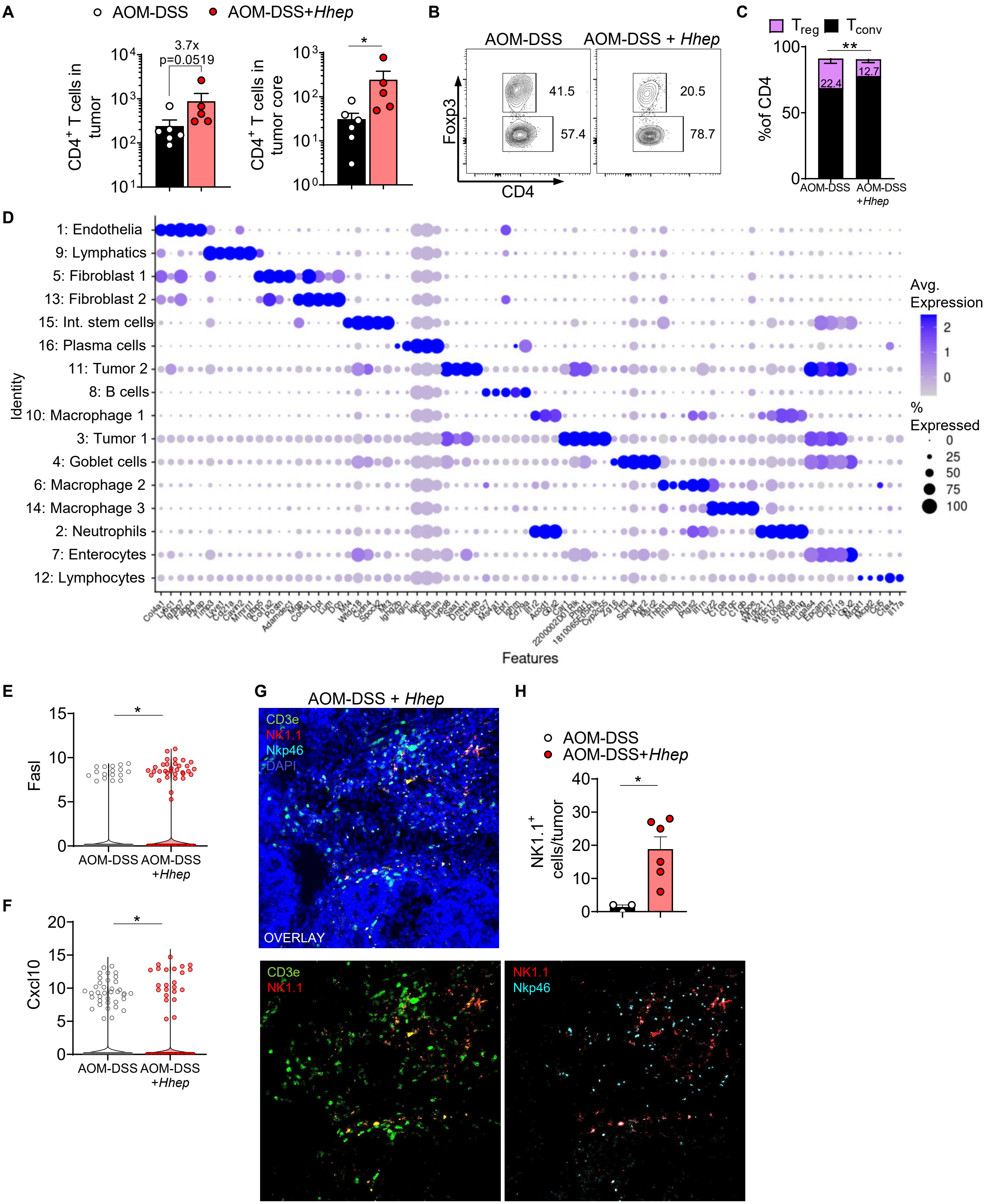

### Supplemental Figure 2: Colonization leads to an increase in cytotoxic lymphocytes within the epithelial layer.

**Figure S2.** *C57Bl/6 Hhep*-free mice were injected i.p. with 10mg/kg AOM on D0 and given 3% DSS in their drinking water on D7-14, 28-35, and 49-56. Half of the mice were gavaged on D45 and 49 with *Hhep.*

(**A**) Quantification of CD4 T cells within the total tumor or the tumor core.

(**B**) Cells were isolated from the tumors of mice +/- *Hhep* 12 weeks post AOM and stained for flow cytometry. Shown are representative graphs of $Foxp3^{+}CD4^{+}$ T cells.

(**C**) Quantification of (**B**).

(**D**) Cells from LP and EL at week 12 of the AOM-DSS protocol (3 AOM-DSS mice, 3 AOM-DSS+Hhep mice) were enriched for $CD45^{+}$ cells, labeled with CD45/MHCi cell hashing antibodies, and subjected to scRNAseq. Bubble plot visualization of differential genes in each cluster. Top differential genes driving clustering in scRNAseq samples. 16 unique clusters were identified from the LP and EL.

(**E-F**) Violin plots of various genes found within the EL of AOM-DSS+/- Hhep mice.

(**G-H**) Colons were sectioned and stained for immunofluorescence 9 weeks post AOM injection.

(**G**) Representative image of staining. Colors shown: CD3e (green), NK1.1 (red), Nkp46 (cyan), and DAPI (blue).

(**H**) Quantification of (**G**).

Data are a composite of 2-3 (**A-C, E-H**) or 1 (**D**) independent experiments with 3-5 mice per group per experiment. Error bars represent the mean ± SEM. Mann Whitney test (**A**), student's T test (**C**), and Wilcoxon rank sum test (**E-F**) were used. $^{*}p<0.05$, $^{**}p<0.005$.

### Supplemental Figure 3: *Hhep* drives a CD4+ $T_{FH}$ response in CRC.

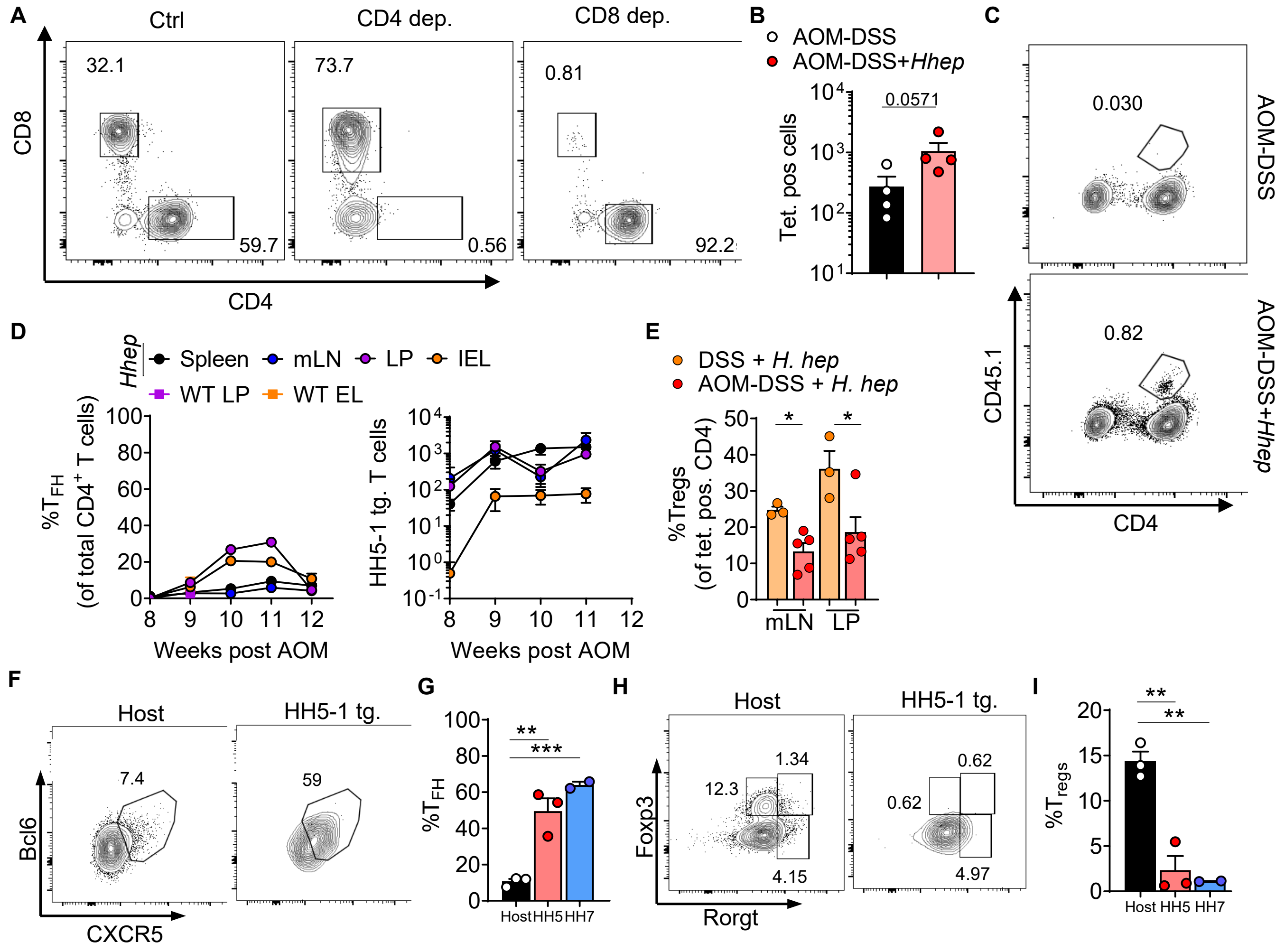

Supplemental Figure 3: *Hhep* drives a CD4$^+$ $T_{FH}$ response in CRC.

**Figure S3.**
(**A**) *C57Bl/6 Hhep*-free mice were injected i.p. with 10mg/kg AOM on D0 and given 3% DSS in their drinking water on D7-14, 28-35, and 49-56. Half of the mice were gavaged on D45 and 49 with *Hhep.* Mice were given anti-CD4 or anti-CD8 every 3 days beginning on day 58. Cells were isolated from the mLN prior to staining for flow cytometry. Representative staining after CD4 or CD8 depletion 12 weeks post AOM.
(**B-F**) Cells were isolated from the colons of tumor-bearing mice +/- Hhep 12 weeks post AOM and enriched for CD45.1$^+$ or Tetramer$^+$ cells prior to staining.
(**B**) Quantification of flow staining of HH-E2 tetramer and CD44 to identify tetramer positive CD4 T cells from the LP of tumor-bearing mice week 12 post AOM +/- *Hhep* colonization at week 7.
(**C**) Cells were isolated from the tumors of mice +/- *Hhep* 12 weeks post AOM and stained for flow cytometry. Shown are representative flow staining of CD45.1$^+$ HH5-1 tg CD4$^+$ T cells. (**D**) Time course of the percent of $T_{FH}$ of total CD4$^+$ and total number of HH5-1tg T cells from different tissues, as indicated.
(**D**) Percentage of $T_{regs}$ within the tetramer positive fraction of cells of *Hhep*-colonized mice treated with either AOM-DSS or DSS alone.
(**E**) Percent $T_{regs}$ of tetramer$^+$ CD4$^+$ T cells after AOM-DSS+*Hhep* or DSS+*Hhep*.
(**F-I**) Cells were isolated from the tumors of mice +/- *Hhep* 12 weeks post AOM and stained for flow cytometry. Percent $T_{FH}$ or $T_{regs}$ of host (black), HH5-1 tg. (red), or HH7-2 tg. (blue) *Hhep*-specific CD4$^+$ T cells in the LP of mice 12 weeks post AOM +/- *Hhep* colonization at week 7.
(**F, H**) Representative flow staining.
(**G, I**) Quantification of (**F, H**).
Data represent 2 (**A,D**) or 3 (**B-C, E-I**) independent experiments with 3-5 mice per group per experiment. Error bars represent the mean ± SEM. Mann Whitney test (**B**) and student's t test (**E, G, I**) were used. *p<0.05, **p<0.005, ***p<0.0005.

Supplemental Figure 4: Colonization leads to an expansion of lymphatics in the LP.

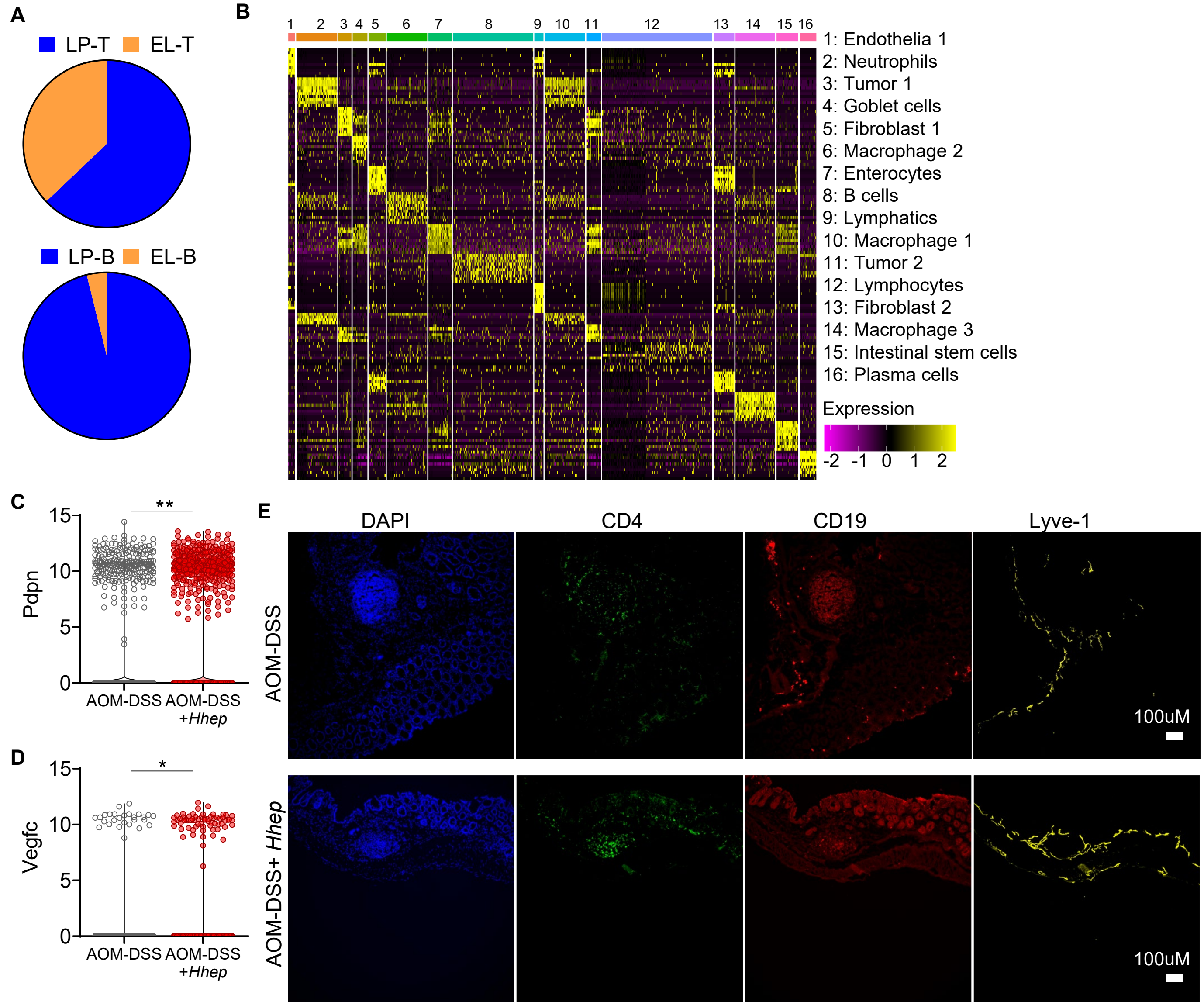


**Figure S4.** Cells from LP and EL at week 12 of the AOM-DSS protocol (3 AOM-DSS mice, 3 AOM-DSS+*Hhep* mice) were enriched for CD45$^{+}$ cells, labeled with CD45/MHCi cell hashing antibodies, and subjected to scRNAseq.

(**A**) Percentage of T and B cells found in the LP and EL fractions from scRNAseq.

(**B**) Top differential genes driving clustering in scRNAseq LP samples. 16 unique clusters were identified. Heatmap visualization of differential genes in each cluster.

(**C-D**) Violin plots of various genes within the LP from scRNAseq.

(**E**) Individual channel breakdowns of Lyve1$^{+}$ IF staining from the LP of mice 12 weeks post AOM +/- *Hhep*. Shown: CD4 (green), CD19 (red), Lyve1 (yellow), DAPI (blue).

Data represent 1 (**A-D**) or 3 (**E**) independent experiment with 3-4 mice per group per experiment. Wilcoxon rank sum test (**C-D**) was used. *p<0.05, **p<0.005.

Supplemental Figure 5: *Hhep* increases peri- and intratumoral TLS.

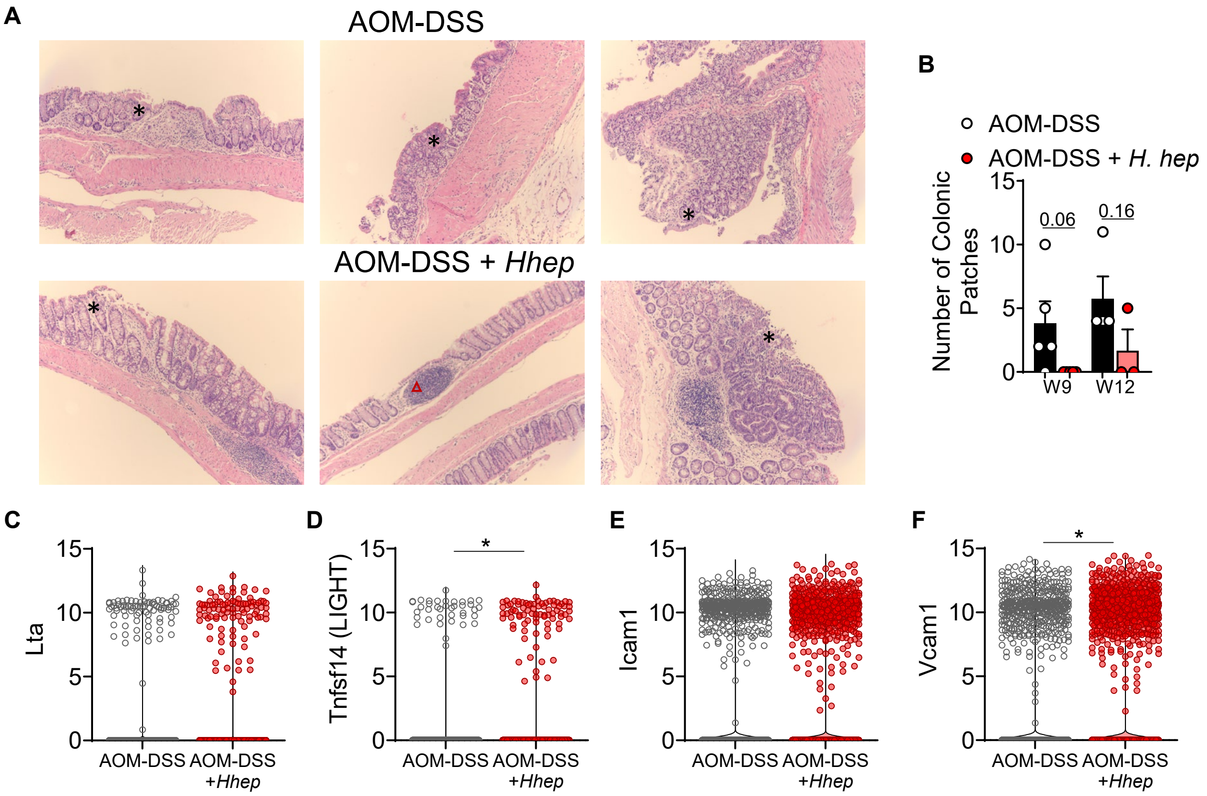


**Figure S5.**

(**A**) H&E sections of the colon of AOM-DSS+/- *Hhep* mice 12 weeks post AOM treatment, highlighting probable tumor tissue (asterisks) and TLS (red triangles).

(**B**) Colons were harvested 12 weeks post AOM and stained for CD4 (green), CD19 (red), CD11c (purple), and DAPI (blue) to identify TLS and colonic patches. Quantification of colonic patches at week 9 and 12.

(**C-F**) Violin plots of various genes within the LP from scRNAseq.

Data represent 1 (**C-F**), 2 (**A**) or 3 (**B**) independent experiments with 3-5 mice per group per experiment. Error bars represent the mean ± SEM. One way ANOVA (**B**) and Wilcoxon rank sum test (**C-F**) was used. *p<0.05.

Supplemental Figure 6: Colonized mice develop *Hhep* containing TLS.

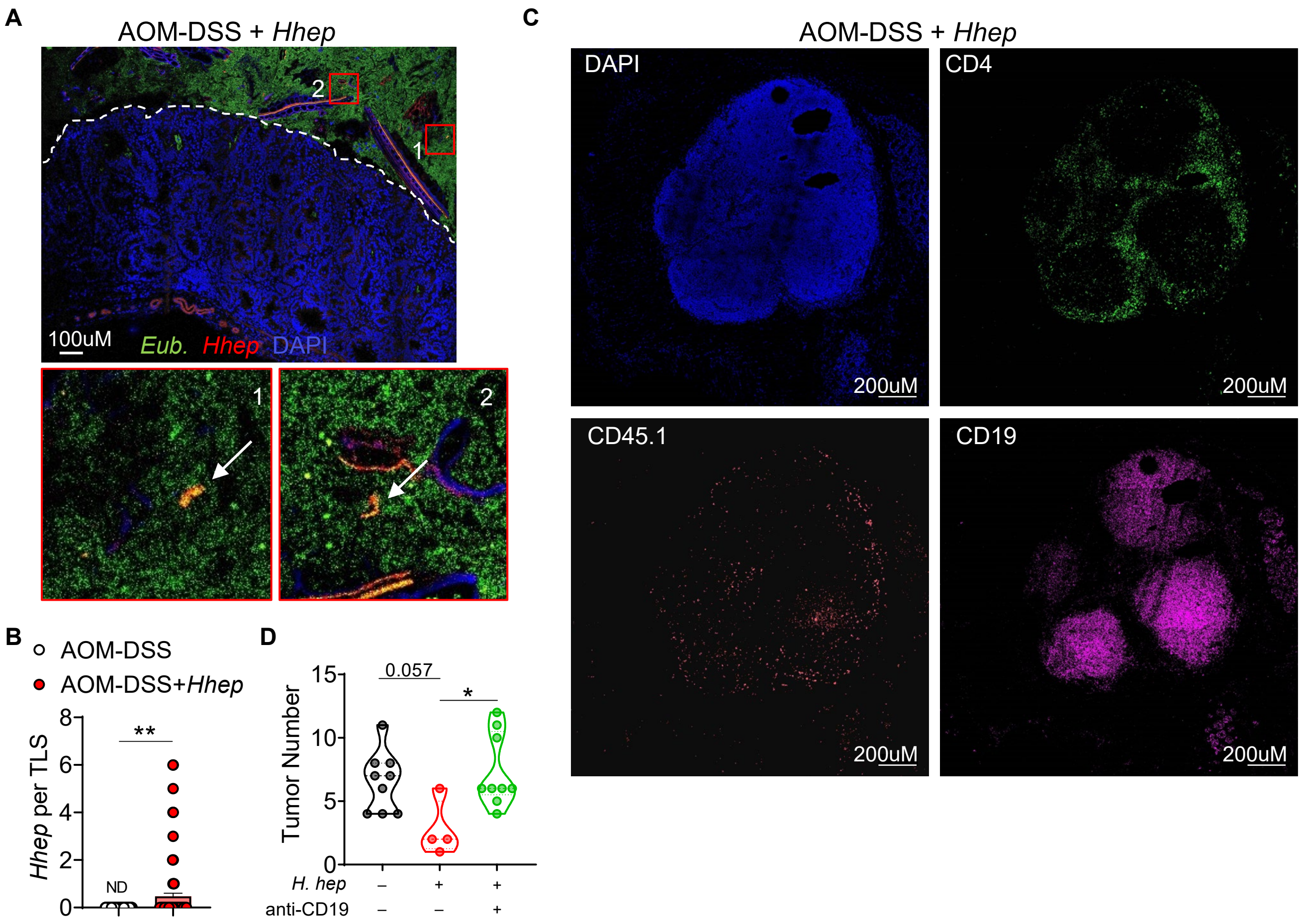


**Figure S6.**

(**A**) Colons were harvested 12 weeks post AOM and fixed in methacarn prior to staining with 16S *Eubacteria* (green), *Hhep* (red) probes and DAPI (blue). Shown are representative images of the colonic lumen.

(**B**) Quantification of *Hhep* per TLS.

(**C**) Single channel breakdown of DAPI, CD4, CD45.1 (HH5-1tg), and CD19 within TLS of an AOM-DSS + *Hhep* mouse.

(**D**) Tumor number quantification from mice given AOM-DSS +/- *Hhep* where half the mice were treated with anti-CD19 every 5 days starting on D58.

Data represent 3 (**A, C-D**) or 2 (**B**) independent experiments with 3-5 mice per group per experiment. Error bars represent the mean ± SEM. Student's t tests (**B**) and one-way ANOVA (**D**) were used. *$p<0.05$, **$p<0.005$.

Supplemental Figure 7: *Hhep*-specific $T_{FH}$ are required for immune invasion into the tumor.

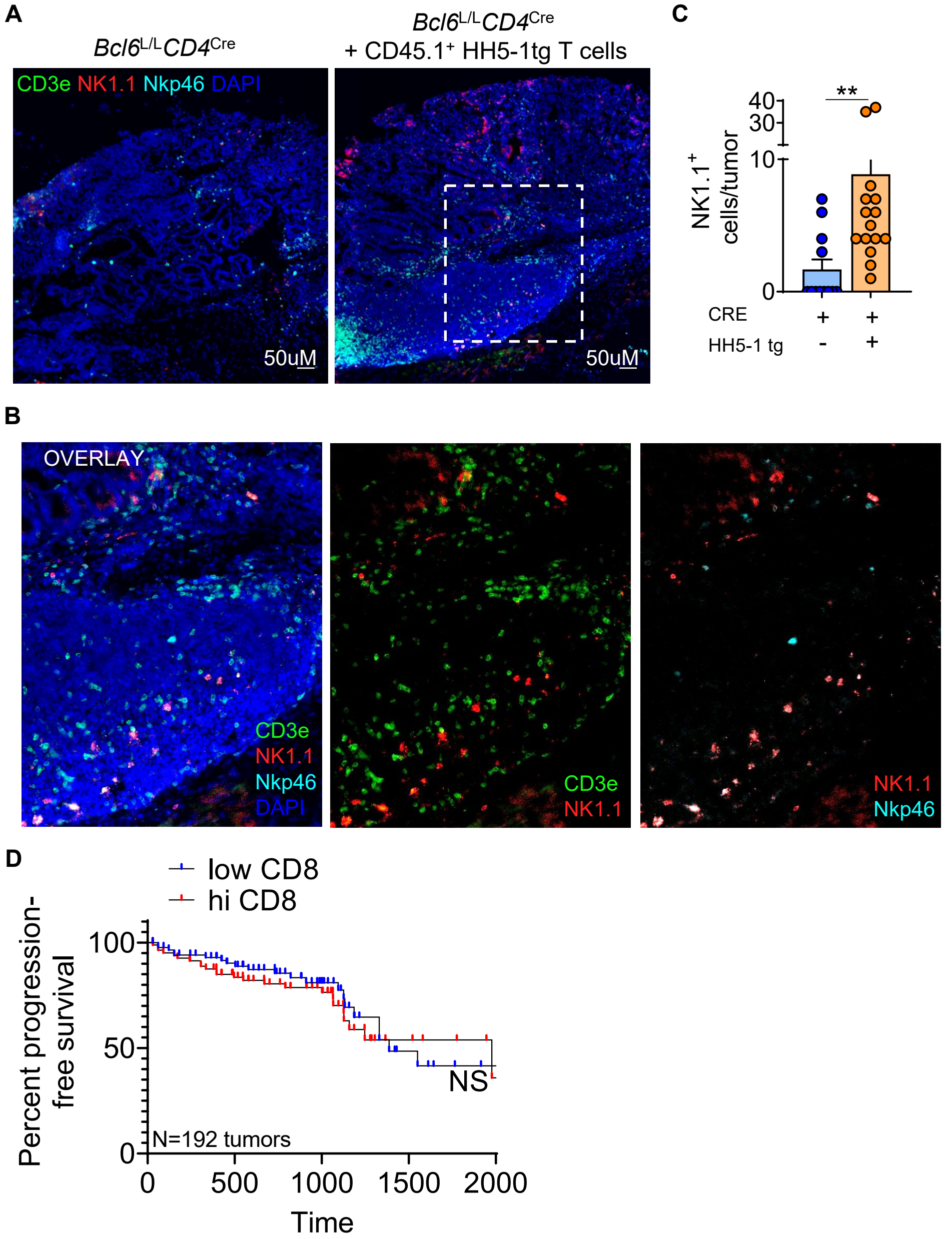


**Figure S7.**

(**A-F**) *Bcl6*$^{L/L}$*Cd4*$^{Cre}$ mice were treated with AOM-DSS + *Hhep +/-* CD45.1$^{+}$HH5-1tg T cells and immune infiltration and TLS numbers were assessed at week 12.

(**A**) Colon sections were stained for CD3e (green), NK1.1 (red), Nkp46 (cyan), and DAPI (blue) to assess infiltration. (**B**) Overlays of all channels, CD3e/NK1.1, and NK1.1/Nkp46.

(**C**) Quantification of (**B**).

(**D**) Progression-Free Survival analysis of patients with hi or low CD8 signatures.

Data are a composite of 2 independent experiments with 3-5 mice per group (**A-C**)**.** Error bars represent the mean ± SEM. Mann Whitney test (**C**) was used. **p<0.005.

#### STAR Methods

#### Experimental Models and Subject Details

Animal Models

6-week old C57Bl/6 mice from Jackson Labs were used for the majority of the studies. *Bcl6*$^{L/L}$*Cd4*$^{Cre}$ mice were acquired from the Poholek Lab, and *Hhep.tg* mice were purchased from Jackson Labs. Both males and females were used and randomly assigned to experimental groups. All mice were maintained at and all experiments were performed in an American Association for the Accreditation of Laboratory Animal Care-accredited animal facility at the University of Pittsburgh and housed in accordance with the procedures outlined in the Guide for the Care and Use of Laboratory Animals under an animal study proposal approved by the Institutional Animal Care and Use Committee of the University of Pittsburgh. Mice were housed in specific pathogen-free (SPF) conditions.

Microbe Strains

*Helicobacter hepaticus* (*Hhep*) 51449 was purchased from ATCC and grown in an anaerobic chamber in either supplemented TSB broth or on chocolate agar plates.

#### Method Detail

Disease models

To establish carcinogen-induced colorectal cancer in mice, we injected 10mg/kg Azoxymethane (AOM) i.p. on D0, prior to 3 cycles of a ‘1 week on, 2 weeks off’ schedule of administering 3% DSS in the drinking water beginning on D7. Mouse weights were monitored weekly and euthanized if more than 25% weight was lost. For *Hhep* colonization, mice were gavaged with ~1x$10^8$ (200uL) two times, 5 days apart at week 7. Mice were sacrificed at either week 9 or 12 post AOM, and tumors were enumerated and measured. For survival experiments, mice were euthanized due to rectal prolapse, weight loss, or general malaise. Attempts to treat minor prolapses were performed for 24hrs, and mice were euthanized if no improvement was made.

*In vivo* treatments

For CD4 or CD8 depleting experiments, mice were treated with 200ug anti-CD4 or anti-CD8 every 3 days beginning week 9 through the end of the experiment. For B cell depletion, mice were treated with 300 ug anti-CD19 every 5 days beginning week 9 through the end of the experiment.

For TCR tg cell transfers, spleens were harvested from naïve $CD45.1^+$ *Hhep* TCR tg mice and stained for naïve T cell markers. Naïve CD4 T cells ($CD90^+CD4^+CD25^-CD44^{LO}$) were sorted at $1x10^5$ cells were transferred r.o. per mouse at week 7, 1 day prior to *Hhep* gavage.

Immunofluorescence

*Tissue and slide preparation*

Tissues were fixed in 1% PFA for 1 hour at 4 deg, then placed in 30% sucrose at 4 deg overnight or until fully dehydrated. Tissue segments were frozen in OCT over dry ice and stored at -80 deg. Tissues were sectioned using a cryostat and frozen for storage. To stain slides, tissues were outlined with a pap-pen, washed 5 times with PBS, washed 5 times with 0.5% BSA in PBS, blocked with 10% rat serum (in 0.5% BSA/PBS) for 45 minutes at RT, stained with primary antibodies in 0.5% BSA/PBS overnight at 4 deg. Tissues were washed 5 times with 0.5% BSA/PBS prior to staining with secondary antibodies for 1 hour at 4 deg and staining with DAPI/Hoescht for 5 minutes covered at RT. Prolong Antifade Gold was used to seal slides, and images were taken as soon as possible. Stained slides were stored covered at 4 deg.

*Fluorescent in situ hybridization*

For FISH staining, tissues were immediately fixed in a methacarn solution at 4 deg overnight. Tissue cassettes were washed with PBS and put in 70% ethanol prior to paraffin embedding and sectioning. Sections were deparaffinized in xylene prior to 2 successive ethanol washes (95% and 90%) and rehydration with ddH2O. 16S Eubacteria and *Hhep* specific probes were added at 100nM in hybridization buffer (0.9M NaCl, 20mM Tris-HCl pH 7.2, 0.1% SDS) and left in a humidified chamber within a 56 deg incubator overnight. A separate slide was stained with a 'scrambled' probe for control. Slides were washed with prewarmed wash buffer (0.9M NaCl, 20mM Tris-HCl pH 7.2) prior to DAPI/Hoescht staining. Slides were sealed with Prolong Antifade Gold and imaged immediately.

*Tumor core calculations*

For calculation of tumor core, images were loaded into Elements software and masks were overlaid over entire tumor. The core mask was determined using the Erode function and tumors were normalized such that the core was the ~15% of the total area for each tumor regardless of size or group. The tumor periphery mask was calculated by subtracting the core area from the total tumor area.

<u>Flow cytometry</u>

Mesenteric lymph nodes (mLN), spleens, and colon tissues were harvested in 3% complete RPMI. Lymph nodes and spleens were forced through 70uM strainers to obtain a single cell suspension. To isolate lymphocytes from lamina propria (LP) and epithelial layers (IEL), we performed gut preparations as previously described(Oldenhove et al., 2009). Single cell suspensions were stained with live/dead and required surface markers for 10 minutes on ice, fixed for 45 minutes on ice with either the eBio or BD fixation kits, and stained for intracellular markers for 45 minutes on ice. For cytokines, cells were stimulated with PMA/Ionomycin and Brefeldin A for 2.5hrs at 37 deg (T cells) or stimulated with BFA alone for 1.5 hrs at 37 deg (innate cells) prior to staining. For TCR tg transfer or tetramer experiments, mice were sacrificed at either week 9 or week 12, and single cell suspensions were made as described above. For tetramer experiments, cell suspensions were stained with tetramer for 1 hour at RT as previously described(Moon et al., 2007). TCR tg or tetramer cells were enriched through a positive selection magnetic pulldown (Stem Cell Technologies) for either the tetramer or CD45.1 (*2*), and both bound and unbound fractions were run by flow cytometry. All flow cytometry was acquired on an LSRFortessa FACS analyzer and cell sorting was carried out on a FACS Aria (BD Biosciences).

scRNAseq

*Library preparation and sequencing*

Libraries were prepared using 10X 5' v1 single-cell RNAseq kit. Briefly, samples from multiple mice derived from either IEL or LP were multiplexed by staining with cell hashing antibodies (Biolegend TotalSeq-C). Sample were then loaded into 2 lanes on a single-cell Chip A, encapsulated into droplets containing individual cells and beads, and then reverse transcribed. Libraries were prepared for sequencing as per the manufacturer's recommendations. Fragment sizes and concentrations of the final prepared gene expression and cell hashing libraries were quantified by BioAnalyzer, and samples were pooled for sequencing on a NovaSeq6000 S2 flow cell at the UPMC Genome Core with the following read parameters: read 1: 28 cycles; i7 index: 8 cycles; i5 index: 0 cycles; read 2: 91 cycles.

*Alignment and generation of gene barcode matrices*

Following sequencing, raw data was demultiplexed into FASTQ files using bcl2fastq from Illumina. Individual FASTQ files from gene expression libraries were then aligned to the mm10 reference genome using CellRanger v3.1.0 (10X Genomics), resulting in generation of gene barcode matrices. For cell hashing libraries, we utilized CITE-seq-Count (https://hoohm.github.io/CITE-seq-Count/#how-to-cite-cite-seq-count) to generate feature-barcode matrices containing read counts for cell hash antibodies by cell barcode. Individual samples were then identified by unique expression of cell hash antibodies associated with individual samples. Cell containing counts for more than one set of CITEseq antibodies were excluded as doublets.

*Dimensionality reduction, clustering and cell type identification*

After generating filtered gene barcode matrices from individual samples, we utilized the R package Seurat (v3.1.4) for downstream processing in R v3.6.1. Gene barcode matrices were first read into Seurat, and a data integration workflow was utilized to integrate data between the two sample sources (i.e. IEL and LP) as described previously (T Stewart, A Butler et al, Cell 2019). Briefly, for each individual sample, library size was normalized, highly variable features were selected, and gene expression was scaled across all cells. Next, dimensionality was reduced using principal component analysis (PCA) on the scaled data and significant principal components were selected heuristically based on the frequency of variance explained. To integrate data, integration anchors were next identified across samples, and used to normalize expression across datasets. Using the integrated data, we then scaled expression values across all cells in the dataset and performed PCA followed by UMAP to visualize cells in a 2-dimensional space. DeteRministic Annealing Gaussian mixture mOdels for clusteriNg Single-Cell data (DRAGON) was then used on the significant principal components to identify clusters. Differentially expressed genes were then identified using a Wilcoxon rank-sum test across clusters as implemented in Seurat, and cell types were characterized based on expression of canonical lineage markers. Downstream analysis to identify more subtle differences within individual lineages was performed by isolating populations of interest, identifying highly variable genes within the lineage of interest, and repeating the dimensionality reduction and visualization workflow for the entire dataset. Raw and processed RNAseq data are available upon request and will be publicly released following acceptance.

**Quantification and Statistical Analysis**

TCGA data analysis

To assess significance of select immune cell populations in overall and progression-free survival of CRC patients, we utilized RSEM normalized $\log_2$ bulk mRNASeq expression data from the TCGA accessed through the Firehose pipeline hosted by the Broad Institute as previously described(Cillo et al., 2020; Deng et al., 2017). Firebrowse was used to identify patient CRC sample cohorts and to download bulk mRNASeq data. Clinical and outcomes data were accessed through the Pan-Cancer clinical Data(Liu et al., 2018). To determine whether there was a relationship between expression of select immune cell population related gene sets and clinical outcomes in patients, we utilized CIBERSORT(Chen et al., 2018) to deconvolve cell frequencies using LM22. Patient outcomes and survival were integrated with CIBERSORT data to determine association between various immune subsets and progression-free or overall survival as well as tumor staging.

16S data analysis

To assess changes in the microbiota over time, stool was taken from C57Bl/6 experimental mice 4 timepoints throughout tumor progression, beginning at D0 post AOM and ending at D82 when mice were sacrificed. Stool was frozen at -80° until the last samples were acquired. Bacterial DNA was isolated from stool using the Qiagen DNA Stool Mini Kit and quantified using a Nanodrop. PCR amplification of the small subunit ribosomal RNA gene (16S rRNA) was performed as follows: DNA were denatured at 94°C for 3 minutes, amplified at 94°C for 45s, 50°C for 60s and 72°C for 90s to amplify, and held at 72°C for 10 min for a final extension step. Microbiome informatics were performed using QIIME2 2020.2(Bolyen et al., 2019). Samples were sequenced by BGI Genomics. Raw sequences were quality-filtered and denoised with DADA2(Callahan et al., 2016). Amplicon variant sequences (ASVs) were aligned with mafft and used to construct a phylogeny with fasttree2(Katoh et al., 2002; Price et al., 2010). Alpha diversity metrics (observed OTUs), beta diversity metrics (Bray Curtis dissimilarity) and Principle Coordinate Analysis (PCoA) were estimated after samples were rarefied to 63,000 (subsampled without replacement) sequences per samples. Taxonomy was assigned to ASVs using naive Bayes taxonomy classifier against the Greengenes 18_8 99% OTUs reference sequences(McDonald et al., 2012). LEfSe was used to compare family level relative abundances between groups. All plots were made with publicly available R packages.

**Statistics**

Data are presented as mean ± SEM. Statistical significance was determined using unpaired Student's t test when comparing two groups, and one-way ANOVA with multiple comparisons, when comparing multiple groups. Some experiments comparing non-parametric groupings (numbers of cells) were compared with a Mann-Whitney test. All statistical analysis was calculated using Prism software (GraphPad). For details on significance, please see figure legends.

**Key Resources Table**

| REAGENT or RESOURCE | SOURCE | IDENTIFIER |
|---|---|---|
| **Antibodies** | | |
| CD4 (depleting) | BioXCell | BE0003-1,<br>RRID:AB_1107636 |
| CD8 (depleting) | BioXCell | BE0117,<br>RRID:AB_10950145 |
| CD19 (depleting) | BioXCell | BE0150,<br>RRID:AB_10949187 |
| CD4 | Fisher | 563727 |
| CD8 | eBio | 11-0083-82 |
| CD19 | Fisher | 562701 |
| Lyve-1 | ReliaTech | 103-PA50 |
| CD11c | Fisher | BDB560584 |
| CD45.1 | eBio | 47-0453-82 |
| PD1 | Biolegend | 135225 |
| CXCR5 | Biolegend | 145532 |
| Bcl6 | BD | 562401 |
| Foxp3 | Fisher | 53-5773-82 |
| Rorgt | eBio | 50245520 |
| CD11b | eBio | 25-0112-82 |
| CD103 | Fisher | 557494 |
| *Hhep* E2 tetramer | NIH | H-$2^b$<br>QESPRIAAAYTIKGA |
| **Bacterial and Virus Strains** | | |

| | | |
|---|---|---|
| *Helicobacter hepaticus (Hhep)* | ATCC | Fox et al. 51449 |
| **Biological Samples** | | |
| **Chemicals, Peptides, and Recombinant Proteins** | | |
| AOM | Sigma | A5486-25mg |
| DSS | MPBio | 16011090 |
| **Critical Commercial Assays** | | |
| **Deposited Data** | | |
| scRNASeq Raw Data | This paper | Available upon request. |
| 16S Raw Data | This paper | BioProject ID: PRJNA655517 |
| **Experimental Models: Cell Lines** | | |
| **Experimental Models: Organisms/Strains** | | |
| $Bcl6^{L/L}Cd4^{Cre}$ | Poholek Lab | |
| *HH5-1 tg* | Jackson Labs | Stock: 032539 |
| *HH7-2 tg* | Jackson Labs | Stock: 032538 |
| **Oligonucleotides** | | |
| *Hhep* FISH probe AF594 | | CCCACACTCCAGAGT AACAGT |
| *Eub.* FISH probe AF647 | | GCTGCCTCCCGTAGG AGT |
| Scramble FISH probe | | CGTGCGGGCATTGCC CTA |
| **Recombinant DNA** | | |
| **Software and Algorithms** | | |

#### Other
